# The Keratin Cortex Stabilizes Cells at High Strains

**DOI:** 10.1101/2025.02.24.639846

**Authors:** Ruth Meyer, Ulla Unkelbach, Parul Jain, Ulrike Rölleke, Nicole Schwarz, Amaury Perez-Tirado, Anna V. Schepers, Claudia Geisler, Andreas Janshoff, Sarah Köster

## Abstract

The eukaryotic cytoskeleton consists of three filament types: actin filaments, microtubules and intermediate filaments (IFs). IF proteins are expressed in a cell-type specific manner, and keratins are found in epithelial cells. In certain cell types, keratin forms a layer close to the membrane which may be referred to as an “IF-cortex”. It is hypothesized that this IF-cortex arranges with radial bundles in a “rim-and-spokes” structure in epithelia. Based on this hypothesis, IFs and actin filaments might add complementary mechanical properties to the cortex. It was previously shown that single IFs in vitro remain undamaged at high strains and display a non-linear stretching behavior. We now ask the question of whether this unique force-extension behavior of single IFs is also relevant in the context of a filament network within a cell. We show that keratin-deficient (KO) MDCK II cells readily form 2D cell layers and 3D cysts and withstand high equibiaxial strains. High-resolution imaging using STED microscopy reveals altered actin cortex structures in KO cells, presumably in response to the missing keratin. We investigate the influence of the equibiaxial strain on the viscoelastic properties of wild-type (WT) and KO cells using atomic force microscopy. We find that the KO cells exhibit a higher pre-stress than the WT cells, likely due to the change of the cortical structure. Interestingly, both the pre-stress and the fluidity of the KO cells are altered already at intermediate strains, whereas the WT cells show a response only at high strain. Similarly, the KO cysts are stretched more easily at low strains than the WT cysts during injection experiments. The compressibility modulus is analyzed in a spatially resolved manner and we find this modulus to be increased at the cell rim, compared to the inside region, due to the geometry of the cell layer. Our results indicate that KO cells compensate for the missing keratin, but are nevertheless very sensitive to external strain, whereas the intricate interplay between the actin and keratin cortices in WT cells preserves the mechanical state and cell stability.

## Introduction

Cells in certain types of tissues permanently experience forces, caused by, for example, fluid flow and active contraction generated by neighboring cells or during peristaltic movement.^1,2^ The cell can cope with these forces due to its cytoskeleton,^1^ a biopolymer network of three types of filamentous proteins – actin filaments, microtubules and intermediate filaments (IFs)^3^ – along with passive cross-linkers and active motor proteins. Interestingly, each type of filament contributes unique mechanic and dynamic properties and together they organize into complex networks providing the cell with distinct mechanical properties and enabling the cell to flexibly adapt to mechanical requirements.^3^ Out of the three filament types, actin is considered to be the most relevant for cell mechanics, ^4^ while microtubules contribute only little.^4,5^ The importance of actin for cell mechanics and morphogenesis is reflected in the so-called cortex, a dense, contractile network found just underneath the plasma membrane. ^6^ In contrast to actin and tubulin, IF proteins are expressed in a cell-type specific manner, each providing their own unique properties to the cell.^7^ Specifically, epithelial cells express the IF protein keratin.^8^ In certain cell types, such as Madin-Darby canine kidney (MDCK) cells,^9^ murine trophectoderm cells or murine enterocytes,^10^ the IF network forms a “rim” close to the cell membrane, just below the actin cortex and may hence also be referred to as an “IF-cortex”. The cortex interconnects the desmosomes in a circumferential network organized in parallel to the plasma membrane and is hypothesized to be part of a rim-and-spokes arrangement.^9^ Furthermore, it has been shown that cells exhibit different mechanical properties in different regions of the cell,^11–13^ which raises the question of whether the rim-and-spokes arrangement brings additional interesting features to the properties of the cell, depending on the region.

In vitro rheology experiments on networks of actin, microtubules, and IFs show that actin and microtubules rupture at comparatively low forces, while IFs are highly extensible and remain undamaged at high strains. ^14–16^ This network extensibility found for IFs is also reflected on the single filament level: vimentin IFs can be extended up to at least 4.5- fold their original length^17^ and similarly high values were found for keratin, desmin, and neurofilaments.^18–20^ Additionally, IFs are referred to as the “safety belt” of the cell as they are easily extensible at slow strain rates, but stiffen when stretched fast. ^17,21–24^ Hence, a “two-layered” cortex in the cells made of mechanically highly distinct filament types, actin and IFs, could potentially lead to interesting properties of the cell when exposed to high forces. Moreover, it has been shown that vimentin IFs stiffen under compression, both in cells and in vitro^25^ and it is not unlikely that a similar phenomenon may exist for keratin.

These results from studies on IF networks and single filaments reveal two interesting features of this biomaterial: (i) they are more extensible than actin filaments that rupture at low strains; and (ii) they display a distinctly non-linear stress-strain behavior. At low strains, IFs behave linearly, at strains above 10 to 20 %, they display a plateau regime in which little force is needed to extend the filament further, and at high strains, they stiffen. ^17^ Thus, the plateau in the stress-strain behavior coincides with a strain-regime where actin is already fluidized^26^ and the question arises of whether the keratin is able to protect the cell from damage. To answer this question, cells may be strained by mechanical stretching^27^ and indeed, such cell stretchers, where cells are seeded on elastic membranes that are being stretched either uniaxially^24,28–31^ or (equi)biaxially^32–34^ have been developed and applied. Alternative methods for stretching of cells exist, such as suspended cell monolayers,^2,35^ single cells between two plates^36^ or cell “domes”.^37^ These studies revealed a strain-stiffening of the cells already at 14% linear strain. ^33^ The actin network reorganizes along the stretching direction,^26,28,30,31^ while IFs reorganize as well but are much more extensible and stabilize the cell.^24,35^

Despite these insightful studies, which clearly point to an important role of IFs in cell mechanics, it remains unclear, whether and to which extent the unique stress-strain behavior of individual IFs is relevant in the cell. We hypothesize that especially at high strains, the high extensibility and the non-linear stress-strain behavior of IFs influence the viscoelastic properties of the cells. To investigate this hypothesis on MDCK II epithelial cells, we employ an equibiaxial cell-stretcher, compatible with both atomic force microscopy (AFM) and light microscopy. As a complementary approach, we present a method to stretch three-dimensional (3D) MDCK II tissue to high strains through microinjection, enabling us to stretch cell layers without the need of an elastic substrate as well as accelerating the stretching process in comparison to the equibiaxial cell stretcher to investigate the safety belt mechanism, i.e. the loading rate dependence of IF mechanics. We study the influence of the keratin network on cell mechanics by conducting AFM force spectroscopy on equibiaxially stretched wild-type (WT) and keratin knock-out (KO) cells. By applying a suitable viscoelastic model to our data, we obtain three viscoelastic parameters, i.e., the pre-stress, fluidity and area compressibility modulus. We perform complementary experiments by stretching 3D cell tissue, so-called cysts, through oil injection. We find that the pre-stress *σ*_0_ is increased in keratin-deficient cells, presumably due to a change of the actin cortex as a compensation of the missing keratin layer. Interestingly, the fluidity value *β* and the pre-stress *σ*_0_ of the KO cells respond to intermediate strains, whereas for the WT cells they respond only for high strains. Similarly, the KO cysts can be stretched more easily than the WT cysts, altogether pointing at a complex interplay between the actin and the keratin cortices in MDCK cells, which is crucial for cell mechanics. Furthermore, we distinguish between the cell edge and interior in a spatially-resolved manner and find that the compressibility modulus *K*^-^*_A_* is increased at the cell rim compared to the inside region as a consequence of the “brick-like” geometry of the cell layers. Together with high-resolution imaging of the actin cortex, our findings suggest that in KO cells actin compensates for the missing keratin network, but only the combination of actin and keratin guarantees cell integrity under strain.

## Materials and Methods

### Cell Stretcher

The home-built stretcher (see Fig. 1(a)) consists of a motorized internal gear ring (dark gray), controlled by the software “spec” (Certified Scientific Software, Cambridge, MA, USA), to which six arms (light gray) are connected via gears. With each motor movement, the gear ring is turned (large pink arrow) resulting in an outwards movement of the arms (small pink arrows) that hold the PDMS device, thereby stretching the device equibiaxially. To ensure unhindered movement and preventing tearing of the device while the arms are turning, the holes in the device are stabilized by poly(tetrafluoroethylene) (PTFE) sleeves, see SI Fig. S1(b). During cell culture and in preparation of the stretching experiments, the PDMS device is kept in a pre-stretcher consisting of a PTFE ring with six pins fitting into the PTFE sleeves in the device and a transfer aid. Thus, the membrane within the PDMS device is kept even and all membranes are set to the same fixed strain at the beginning of the experiments. The transfer aid made from stainless steel is situated between the pre-stretcher and the device and provides a seamless transfer from the Petri dish to the stretcher by ensuring a smooth sliding off of the pins of the pre-stretcher onto the arms of the stretcher.

**Figure 1:**
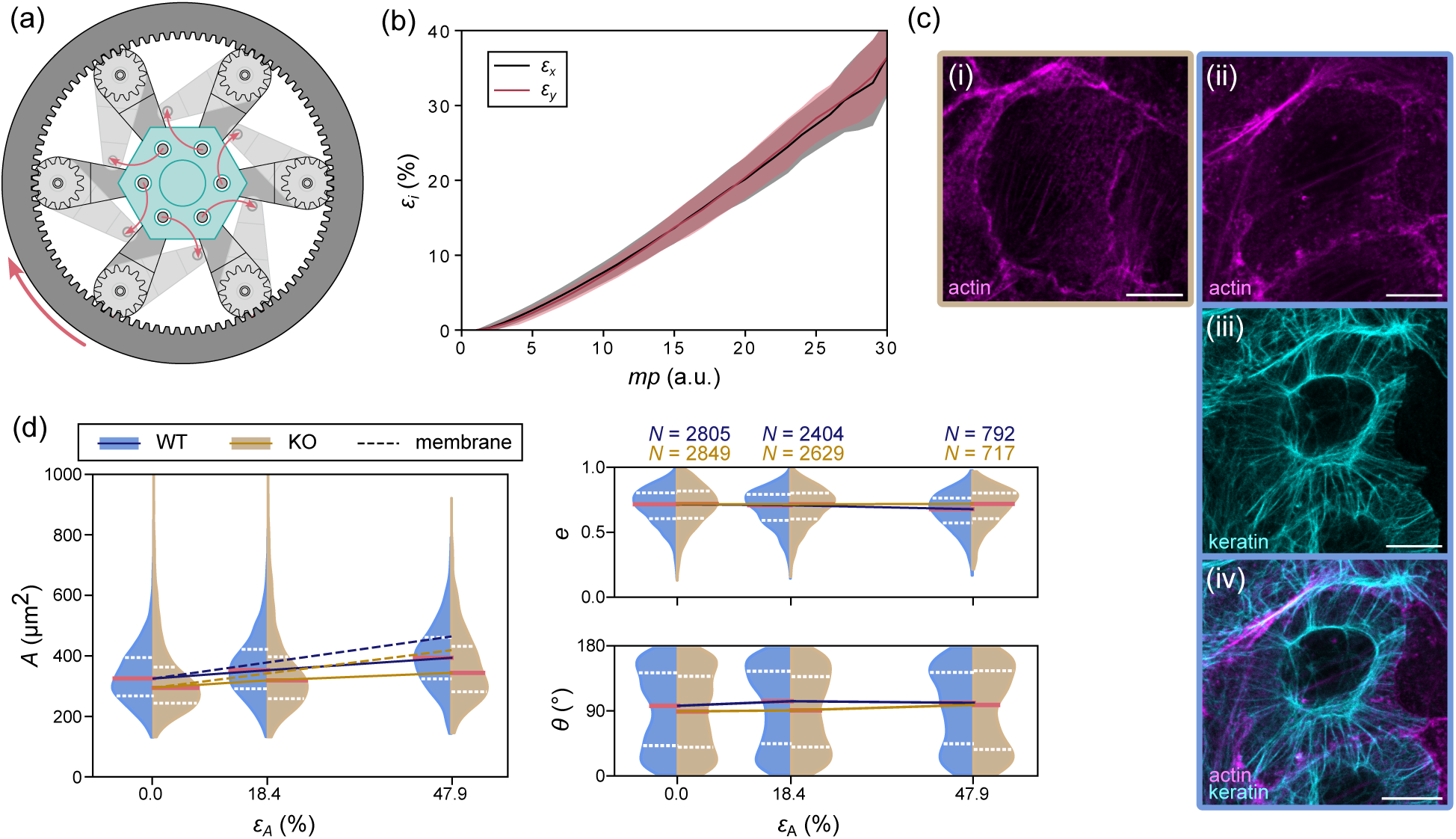
Experimental setup and characterization of the cell stretcher. (a) Schematic of the equibiaxial stretcher. A PDMS device (cyan) is attached to the six arms of the stretcher (light gray). The motor rotates the outer internal gear ring (dark gray, large pink arrow), which then moves the arms outwards (small pink arrows). (b) Averaged linear strain *ε_i_* (*x*- and *y*-direction) of the PDMS membrane, calculated from the beads embedded in the membrane, plotted against the motor steps *mp*; the shaded areas show the standard deviation. (c) Examples of confocal microscopy images of the cell lines used in the study. (i) Fixed, unstrained MDCK II/K8 KO cells with phalloidin-stained actin (magenta), referred to as KO. (ii-iv) Fixed, unstrained MDCK II WT cells transfected with lifeact actin tagged with mCherry, here additionally stained with phalloidin (magenta, ii), and keratin 8 tagged with EGFP (cyan, iii), referred to as WT (composite, iv). Scale bars correspond to 10 *µ*m. (d) Area *A* (left), eccentricity *e* and orientation *θ* (right) of the WT (blue) and KO (yellow) cells plotted against the area strain *ε_A_*. The violin plots show the distribution of the results for each individual cell with the median (solid pink lines) and upper and lower quartiles (dashed white lines). Medians are connected using solid lines. Dashed lines show the theoretically expected area of the cells calculated from the characterization of the membrane. The numbers of cells *N* are provided in the figure and correspond to all data.

### PDMS Devices

We fabricate polydimethylsiloxane (PDMS, Sylgard 184, Dow Chemical, Midland, US) devices with a total diameter of 3.25 cm, see SI Fig. S1(a). They consist of a membrane with a thickness of 250 *µ*m. The walls have a height of 0.5 cm and delimit a well of radius 1 cm on the membrane into which the cells are seeded.

To fabricate the walls of the devices, 3 g of PDMS base are mixed with cross-linker at a ratio of 25:1, filled into home-made hexagonal molds made from polylactic acid (PLA, Polymaker, Shanghai, China), cured for 1.5 h at 65*^◦^*C and removed from the mold. We produce the membranes in two layers. PDMS is mixed with cross-linker at a ratio of 25:1, spin-coated onto a 4-inch Si-wafer (ramp up to 60 rpm within 10 s, 10 s at 60 rpm) to obtain a thickness of 220 *µ*m and cured for 45 min at 65*^◦^*C. The membrane is plasma activated (Zepto, Diener electronic GmbH & Co. KG, Ebhausen, Germany) at 30 W for 21 s. Dark-red fluorescent microspheres (660/680 nm, 2% solids, FluoSpheres F8807, 200 nm, Invitrogen, Waltham, MA, USA) are diluted in a ratio of 1:200 in isopropanol and spin-coated onto the membrane (ramp up to 60 rpm within 25 s, 25 s at 60 rpm). A second layer of PDMS with a mixing ratio of 25:1 is spin-coated onto the same wafer (ramp up to 2460 rpm within, 40 s, 20 s at 2460 rpm) to obtain a thickness of 30 *µ*m.

We place the cured PDMS walls onto the uncured PDMS membrane, cure everything together for 45 min at 65*^◦^*C and remove the device from the wafer. Holes are punched into the wall using a 4 mm biopsy puncher and the PTFE-sleeves are inserted. The finished PDMS devices are boiled in water for 1 h and stored in sterile water for at least 8h. This step ensures that the PDMS device is sterile for cell culture and that the device is soaked with water to prevent the cell medium from being absorbed by the device.

In preparation for the cell experiments, the pre-stretcher is assembled and added to the PDMS device (SI Fig. S1(b)). The well inside the device is rinsed with PBS, coated with laminin (25 *µ*g/mL, from mouse, Gibco, Life Technologies, Carlsbad, CA, US) for 1h at room temperature and rinsed again with PBS.

### Cell Culture

All experiments are performed on Madin-Darby Canine Kidney cells strain II (MDCK II, ECACC 00062107, European Collection of Authenticated Cell Cultures, Salisbury, UK). We stably transfect the cells with a bicistronic vector to allow for coexpression of K8-EGFP and lifeact-mCherry (VectorBuilder, Neu-Isenburg, Germany); this way, we fluorescently tag both the keratin and the actin structures. In the following, we will refer to these cells as wild-type (WT) cells. The keratin 8 knock-out (KO) cells are produced using CRISPR/Cas9n technology. We clone two different gRNAs targeting exon 1 of the canine Krt8 gene into PX462 V2.0 vector (Addgene #62987, Watertown, MA, US; kind gift of Feng Zhang).^38^ The sequences of the gRNAs are gRNA-1: 5’-GGG GCT CAC CTT GTC GAT GAA GG -3’ and gRNA-2: 5’-GGG CGG GCA GTG CCT GGG GCT GG -3’. We co-transfect MDCK II cells with both vectors and subject isolated single cell clones to immunofluorescence analysis using anti-keratin 8 (TROMA-1, DSHB, Iowa City, IA, US) and anti-desmoplakin (DP-1, Progen, Heidelberg, Germany) antibodies and immunoblot analysis to confirm knock-out (SI Fig. S2(c) and (d)).

The cells are cultured in Eagle’s Minimum Essential Medium, supplemented with Earle’s salts and 2 mM GlutaMax (EMEM, Gibco) and 10% fetal bovine serum (FBS, Gibco), and kept at 37*^◦^*C and 5% CO_2_. For the WT cells, we add 2 *µ*g/mL Puromycin (Sigma Aldrich, St. Louis, MO, USA) to the medium. Twice per week, the cells are passaged, and passages 4 to 12 after thawing are used for the experiments.

Two days before the AFM experiments, we seed the cells on the PDMS devices to reach confluence for the experiments. To improve the image quality for fluorescence microscopy, the cell medium is replaced by phenol-free medium (Gibco) containing the same supplements as the regular culturing medium. Additionally, to be able to perform experiments without CO_2_ supply, we add 15 mM HEPES (Thermo Fisher Scientific, Waltham, MA, US) to the cell medium.

To cultivate the cysts, we coat gridded Petri dishes (*µ*-dish, 35mm, low, Ibidi, Gräfelfing, Germany) by incubating them with laminin (20 *µ*g/cm^2^) for 1 h at 37*^◦^*C and 5% CO_2_. For the following steps, all material including the cells are cooled to 4*^◦^*C. 50 000 cells, prepared in 450 *µ*L medium, and 50 *µ*L Matrigel (Corning Inc., Corning, NY, US) are gently mixed. The solution is then transferred into the dish and incubated at 37*^◦^*C and 5% CO_2_ for 7–9 days to allow for cyst formation.

### Immunoblot analysis

Expression of keratin 8 in original, i.e., non-transfected, wild-type and in keratin 8 KO MDCK II cells is analyzed by immunoblotting. Polypeptides are separated by standard SDS polyacrylamide gel electrophoresis and transferred onto polyvinylidene fluoride Immobilon-P membranes (Millipore, Merck, Darmstadt, Germany) by tank blotting. The membranes are blocked with 1x RotiBlock (Roth, Karlsruhe, Germany) and incubated overnight at 4*^◦^*C with rat anti-keratin 8 (TROMA-1, DSHB) and rabbit anti-actin (Sigma Aldrich, A2066) antibodies. Membranes are washed in TBS-T buffer (50 mM Tris, 150 mM NaCl, 0.05% Tween 20, pH 7.6) and subsequently incubated with secondary antibodies (anti-rat HRP, anti-rabbit HRP, both from Dianova, Hamburg, Germany) for 1 h at room temperature. Bound antibodies are detected with SuperSignal West Pico PLUS Chemiluminescent Substrate (Thermo Fisher Scientific) and a chemiluminescence system (Fusion SL, Vilber Lourmat). On the same membranes keratin 8 and actin are detected consecutively. Bound antibodies are stripped by incubation of the membranes with stripping buffer (100 mM glycine, pH 2) three times for 20 min.

### Cell Fixation, Immunofluorescence Staining and Fluorescence Imaging

For the imaging of the actin and keratin networks in both cell lines, the cells are seeded in advance on cover slips (diameter of 18 mm, thickness 1.5H, VWR, Radnor, PA, US). When the cells reach confluence, they are washed with PBS and fixed with 4% formaldehyde (Thermo Fisher Scientific) for 20 min at room temperature. Afterwards, the cells are permeabilized by washing briefly with 0.1% Triton X solution (Triton X-100, Sigma Aldrich, diluted with PBS). To stain the actin, phalloidin (Alexa Fluor 647, Invitrogen) is mixed with 0.1% Triton X solution in a ratio of 1 to 40 and added to the cells for 1 h at room temperature. Then, the cells are washed, first with 0.1% Triton X solution, and second with ultrapure water. Finally, the glass slides are mounted on a microscopy slide with mounting medium (ProLong Diamond antifade, Invitrogen) and stored at 4 *^◦^*C for at least one day.

The fixed and stained cells are imaged on an inverse confocal microscope (IX83 with a Fluoview FV3000, Olympus, Hamburg, Germany) at 100× magnification (NA 1.45, UP-LanXApo, Olympus) and laser line 640 nm for actin and laser line 488 nm for keratin. *Z*-stacks are recorded with Δ*z* = 0.41 *µ*m.

For the high-resolution imaging of the actin networks in both cell lines, cysts are seeded in advance in Matrigel in Petri dishes (*µ*-dish, 35mm, high, Ibidi) to reach confluence and allow for cyst formation. To remove the Matrigel from the cysts, Cell Recovery Solution (Corning Inc.) is added and incubated at 4 *^◦^*C for 25 min. Cysts are washed with PBS and 1 *µ*M staining solution (actin LIVE RED, abberior, Göttingen, Germany) is added to the sample and incubated for 60 min at 37*^◦^*C and 5% CO_2_. Afterwards, the cysts are washed with PBS and 4% formaldehyde (Thermo Fisher Scientific, Waltham, MA, US) is added and incubated for 20 min at room temperature to fix the cells. Finally, the Petri dish is filled with PBS and stored at 4 *^◦^*C until imaging.

STED imaging of actin networks in cysts is performed using a STEDYcon system (Abberior Instruments, Göttingen, Germany) mounted on an inverse fluorescence microscope (IX83, Olympus, Hamburg, Germany) with a 60× water immersion objective (NA 1.20, UPLanSApo, Olympus). The objective’s correction collar is adjusted to account for the coverslip thickness of the Petri dishes used, thereby minimizing spherical aberrations. A 640 nm laser and a 775 nm laser are used for excitation and depletion, respectively, to acquire STED images with increased resolution in the lateral dimension compared to confocal images.

Epi-fluorescence imaging of the fluorescent beads for the characterization of the PDMS stretching devices is performed on a motorized upright fluorescence microscope (BX63, Olympus) at 60× magnification using a water dipping objective (NA 1, LUMPLFLN60XW, Olympus) with an Orca Flash 4.0 camera (Hamamatsu, Tokio, Japan) and a Sola SE II fluorescence lamp (Lumencor, USA) set to 15% power. We image the fluorescence microspheres using a Cy5 filter set (U-F49006, excitation: ET620/60x, dichroic mirror: T660LPXR, emission: ET700/75m; Olympus) at 50 ms exposure time. At each position, *z*-stacks are recorded with Δ*z* = 0.54 *µ*m.

### Characterization of the Cell Stretcher and PDMS Device

Characterization of the stretcher and PDMS stretching devices is performed without cells. The devices are filled with water. The device is stretched in motor steps of Δ*mp* = 0.1 a.u. until rupture.

In the stretcher, the membrane tends to be uneven or tilted because it is manually placed on the stretcher arms (SI Fig. S14(a)). To overcome this problem, we record 3D epi-fluorescence images of the beads and correct the images for the tilt. As the beads are in one layer in the PDMS membrane only and the bead size is smaller than the difference in *z* between two images, for each lateral position there is only one *z*-position where the beads are in focus. We compute a spatial Laplacian for every image for a kernel size of 4 pixels to define the sharpness. We normalize the Laplacian along the *z*-axis with respect to the maximum absolute value in a pixel-wise manner. Next, we determine which slice in the stack is in focus for a coarse grid of the image. The sum of the normalized, absolute gradient values is computed for each *z*-slice and grid block. A 2D matrix is then created storing the indices of the *z*-slice with the highest sum, representing the “best-in-focus” slices at each grid block (SI Fig. S14(b)). Finally, we fit a second order 2D polynomial to the grid and interpolate the original *z*-stack to the result of this fit to create a tilt-corrected image. We repeat this procedure for every stack at every stretching position in the experiment to generate a time series (SI Fig. S14(c)).

For the calculation of the strain, we manually align the tilt-corrected images at all strains in the center (SI Fig. S14(d)). Using the python implementation “itk-elastix” of the software “Elastix”,^39,40^ every image in the image series of the beads is registered with respect to its previous image using an affine transformation (SI Fig. S14(e)).

This procedure is repeated for twenty independent experiments. The arithmetic mean and standard deviation of all data sets is calculated and the values for the linear strain in *x* direction from the averaging for each motor position *mp* is used to determine the area strain (Eq. 4), since *ε_x_* = *ε_y_* applies for equibiaxial strain.

### Atomic Force Microscopy

The force spectroscopy measurements are performed on a Nanowizard 4 atomic force microscope (AFM, Bruker Nano GmbH, MA, USA) combined with a manual or motorized inverse fluorescence microscope (IX73 or IX83, Olympus/Evident, Hamburg, Germany). Three different strain positions (0%, 18.4% and 47.8% area strain) are included for the measurements and at each strain approximately the same region within the device is chosen. We use a cantilever with a spherical tip of 2 *µ*m diameter (CP-PNPL-SiO-A, Nano and More GmbH, Wetzlar, Germany). The maximum force, i.e., the setpoint, is set to 2 nN and we use an indentation velocity of 2 *µ*m/s. Force maps of size 50 *µ*m × 50 *µ*m with a pixel size of 2 × 2 *µ*m^2^ are recorded. At each strain position, we take four force maps to cover a total area of 100 *µ*m × 100 *µ*m. The measurements are performed in liquid in a heated environment at 37*^◦^*C (The Cube, Life Imaging Services, Basel, Switzerland). Phase contrast images are taken at 20× magnification (NA 0.45, LUCPlanFLN, Olympus) with a monochromatic camera (UI-3060CP-M-GL Rev. 2, IDS Imaging Development Systems GmbH, Obersulm, Germany) and 50 ms exposure time. The experiments are terminated either after reaching the highest strain or when the PDMS device breaks.

### Cyst Injection

For the injection experiments, the outer edge of the Matrigel in the Petri dish is removed using a pipette and subsequently, the cysts are washed with PBS at 4*^◦^*C. Cell Recovery Solution (Corning Inc.) is added to the cysts and incubated at 4 *^◦^*C for 25 min. Finally, the sample is washed with PBS and warm cell medium with HEPES (15 mM, Thermo Fisher Scientific) and penicillin/streptomycin (10000 units/(10 mg/mL), Merck, Darmstadt, Germany) is added.

The experiments are performed on a motorized inverse microscope (IX81, Olympus) at 40× magnification (NA 0.95, UPLSAPO40X2, Olympus) and an XM10 camera (Olympus). To carry out the injection of the cysts, a microinjector (InjectMan NI2, Eppendorf, Hamburg, Germany) is used in combination with a micropipette with an inner diameter of 0.5 *µ*m and an outer diameter of 1 *µ*m (Femtotip, Eppendorf) and mineral oil for molecular biology (*ρ* = 0.84 g/mL, BioReagent, Sigma Aldrich). Before the injection, a phase contrast image of the cyst is taken. The tip is inserted into the cyst and oil is released into the lumen of the cyst at an injection pressure varying between 1000–6000 hPa for 0.5–6 s. The amount of oil injected into the cyst is determined by visually inspecting the cyst while injecting in order to reach a sufficient degree of strain, but avoid a rupture of the cyst. Once the oil droplet is big enough, the tip is pulled out of the cyst. During the whole injection process, we record a phase contrast video. In the end, a phase contrast image is recorded as before the injection. Within one Petri dish, up to four cysts are injected. In total, the time between removing the Matrigel and finally fixing the sample is at maximum 1 h. We repeat this procedure on 27 WT cysts and 28 KO cysts from 10 independent experiments each.

### Image Analysis

To analyze the area of the cells, cell segmentation is performed on the phase contrast microscopy images using the software “CellPose” ^41^ and its python implementation (SI Fig. S3(a), (b)). We use the model “cyto2” ^42^ and the default flow threshold of 0.4 for all data sets; here, one data sets corresponds to the data collected from one experiment. 11 data sets are used for the WT cells and 10 for the KO cells. The cell area is determined by summing over the number of pixels per cell and scaling with the pixel size of the images (SI Fig. S3(c)).

For the determination of the eccentricity and orientation of the cells, ellipses are fitted to the outline determined from CellPose using the python function “fitEllipse” in the package “cv2” (SI Fig. S3(d)). The eccentricity of the ellipse is calculated via its semi-major axis *a* and its semi-minor axis *b* by

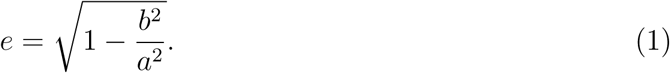

The orientation is defined as the angle between the semi-major axis *a* and the *x*-axis from 0*^◦^* to 180*^◦^* as the direction is irrelevant (SI Fig. S3(e)).

To analyze the cyst shape in the cyst injection experiments over time, we use the video taken during the injection. The workflow is schematically shown in Supplementary Fig. S4. We manually segment the cyst, the lumen and the oil droplet in each individual image frame of the video (SI Fig. S8(a)) using the polygon tool in Fiji^43^ to outline each part and convert the outline into a mask (SI Fig. S8(b)). We calculate the center of mass of the lumen (SI Fig. S8c), determine vectors **r** from the center of mass to the outline of the mask for 200 varying angles *ϑ* from −*π < ϑ <* +*π* and calculate the length of each vector (SI Fig. S8(d)).

Subsequently, the mean of all vector lengths is calculated as the radius of the lumen *r*_Lumen_.

We repeat the process to determine the radius of the cysts *r*_Cyst_ with the same center of mass as the lumen. As the oil droplet is not necessarily positioned in the exact center of the lumen, the center of mass is calculated for the oil droplet separately (SI Fig. S8(c)) and the radius *r*_Oil_ is determined in the same manner as for the lumen and cyst.

This procedure is repeated for every frame in the video, starting from one frame before the first oil droplet is released into the cyst until the stretching is finished, but before the micropipette is pulled out of the cyst (SI Fig. S8(e)).

Finally, the volumes of cyst, lumen and oil droplet are determined with the assumption of a perfect sphere for each and the average thickness *d* of the shell is calculated by

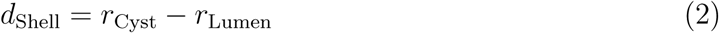

### AFM Data Analysis

For the analysis of the force spectroscopy measurements, we preprocess the data curves using the python package “nanite” ^44^ (SI Fig. S15). At first, a baseline correction is applied to the raw force curves (SI Fig. S15(a)), correcting its tilt and offset to the *x*-axis (b). Next, the contact point is estimated using the nanite-function “deviation from baseline” (SI Fig. S15(c)). Finally, the curve is corrected by the tip-sample separation, which separates the distance of the cantilever towards the sample from the deflection of the cantilever in the opposite direction (SI Fig. S15(d)).

The workflow for the analysis is shown schematically in SI Fig. S4. We combine the data from the force spectroscopy measurements with the images from light microscopy. The phase contrast images are segmented with CellPose as described before (SI Fig. S4(a), (b)). As the phase contrast images have a different pixel size than the AFM images, the resulting cell outlines are interpolated to fit the AFM pixel size (SI Fig. S4(c)). These outlines are then overlaid onto the AFM images where each pixel contains a force spectroscopy curve (SI Fig. S4(d), (e)). Each cell is divided into two regions using the outline. The outline traces the outermost pixels that are still part of the cell. The rim is defined as a single pixel-wide line that marks this edge of the cell. The inside region is then identified as the area in the cell center extending three pixels inward from the rim (SI Fig. S4(f)). With this division, we exclude an interim region between the rim and the inside to avoid any unclear classification into either of these regions.

Finally, the curves are rated using the respective method implemented in nanite. We train our own neural network using the respective feature in nanite on force curves that are not included here in the further analysis to fit with our data and results. We exclude every curve with a rating lower than 4.5 to omit out any curves of low quality (SI Fig. S4(g)).

The force curves are averaged per category by aligning all curves on their maximum. Since the time points of the curves can be different, we interpolate the curves to correct the time axis. Finally, the mean and the standard deviation of the curves are calculated.

The individual curves are fitted with a viscoelastic fit providing the viscoelastic properties prestress of the cortex, apparent compressibility modulus and fluidity. After fitting each curve, we exclude those curves where the viscoelastic fit provides unphysical values and show a failed fitting of the curve.

## Results and Discussion

### Keratin KO cells withstand equibiaxial strain

To investigate cells under strain, we design a cell stretcher (Fig. 1(a)) that pulls on an elastic polydimethylsiloxane (PDMS) membrane (cyan) equibiaxially by turning the six arms (light gray) outwards in an “iris-like” motion (small pink arrows). We first thoroughly characterize the stretcher by adding a layer of fluorescent beads as fiducial markers to the membrane (SI Fig. S1(a)) and by imaging the beads at different motor positions of the stretcher. We analyze the displacement of the beads by registering the images using the software “Elastix” ^39,40^ and thus obtain the elongation of the membrane Δ*L* between two motor positions of the stretcher, which we use to calculate the strain in *x* and *y* direction as

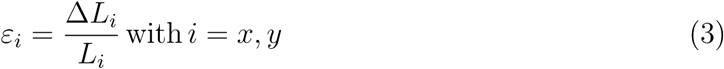

with the initial length *L*.

Fig. 1(b) shows the linear strain *ε_i_* (*i* = *x*, *y*) in *x*- and *y*-direction against the applied steps of the motor *mp* (*N* = 21 independent experiments). The solid lines represent the average from all experiments in *x* (black) and *y* (pink) direction. The shaded areas present the standard deviations. We find the same linear strain in *x* and in *y* direction, confirming that the stretcher indeed pulls uniformly in both directions. This is in contrast to uniaxial strain, where the elastic membrane and therefore the cell sample is stretched in one direction and compressed perpendicularly to the stretching direction according to the Poisson’s ratio.^45^ A good measure to quantify the degree of stretching is therefore the area strain *ε_A_*. The 2-dimensional area strain can be described as

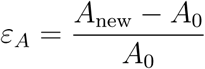

and hence^46^

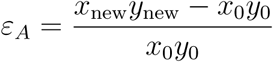

for the area *A* = *x* · *y*. Substituting *x*_new_ and *y*_new_ by *x*_0_ + Δ*x* and *y*_0_ + Δ*y*, respectively, we obtain

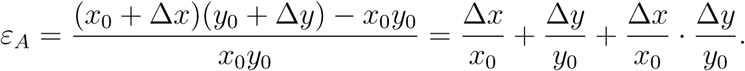

Using the definition of linear strain (Eq. 3) yields

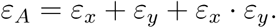

Finally, for equibiaxial strain *ε_x_* = *ε_y_*, the area strain can be written as

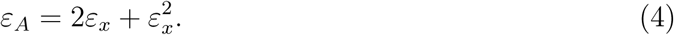

For our experiments, we use epithelial MDCK II cells which naturally express keratin 8/18 in a rim-and-spokes arrangement.^9^ For visualization of the keratin and actin networks, the cells are transfected with keratin 8 tagged with EGFP and lifeact actin tagged with mCherry (Fig. 1(cii)-(civ)). In the following, we will refer to these cells as wild-type (WT) cells. Additionally, to investigate the influence of the keratin network on the mechanical properties, we use MDCK II K8 knock-out (KO) cells (Fig. 1(ci), SI Fig. S2), referred to as KO cells.

Two days before the mechanical experiments, we seed the cells on elastic PDMS membranes and let them grow to a confluent layer. To confirm the stretching of cells grown on the membranes, we image the cells with phase contrast microscopy at increasing motor positions *mp* of the stretcher (*N* = 10–11 independent experiments). Each field of view corresponds to 300 × 300 *µ*m^2^. The images are segmented using the software “CellPose” ^41,42^ and the area *A* of each individual segmented cell is calculated by counting the number of pixels per cell.

Additionally, we fit an ellipse to the outline of each segmented cell to calculate their orientation *θ* and eccentricity *e* (Eq. 1, SI Fig. S3(d), (e)). An eccentricity of 0 and 1 represent a circle and a parabola, respectively. Any value between 0 and 1 represents an ellipse.

Fig. 1(d) shows the results of this analysis for increasing area strains *ε_A_* as violin plots. The WT cells are displayed in blue, the KO cells in yellow. The solid pink lines represent the median of each distribution, the dashed white lines the quartiles. As a guide to the eye, the medians are connected by solid lines in the respective colors. Additionally, we show the theoretically expected value for the area from Eq. 4, considering a perfect and complete transfer of the strain from the membrane to the cells and using the median from each cell line at 0% strain as the starting value (dashed lines).

We find the KO cells to be smaller than the WT cells, as previously also observed for keratin 8 knock-out hepatocytes.^47^ Additionally, we observe an increase in cell size with increasing strain for both cell lines. The increase is lower than expected from the applied strain even though the cell layer remains intact and no holes in the cell sheet appear in the field-of-view. This deviation from the theoretical area values can have multiple reasons. The elastic membrane is coated with laminin to enhance cell adhesion. This attachment of the cells to the coating and the coating to the membrane can change and be disrupted during stretching leading to a lower strain transmission or an uneven strain distribution.^33,34,48,49^ Furthermore, it has been shown that for treatment of the cells with latrunculin, hence weakening the focal adhesions through the inhibition of actin polymerization, the cell strain decreases considerably, proving the relevance of a good adhesion of the cell to the surface.^33^ Additionally, the cell body is mechanically inhomogeneous, which can also lead to a reduced strain transmission on the cells. ^49^ As expected, the orientation and the eccentricity of the cells do not change with increasing strain, see Fig. 1(d), right, supporting the notion of equibiaxial strain.

With this cell stretcher design, we are able to stretch the elastic membrane up to a linear strain of 38% which translates to an area strain *ε_A_* of 90% (Eq. 4) and is higher than for other equibiaxial cell stretchers.^27,32–34^ Here, we stop stretching the elastic membrane at an area strain of 48 % to ensure long-term measurements without risking yielding of the PDMS membrane. Additionally, we choose an intermediate strain approximately halfway between the maximum chosen strain and the relaxed position. Although the applied strain does not fully transfer to the cells, we are able to stretch both cell lines equibiaxially to an area increase of up to 23%. In the following, the area strain *ε_A_* refers to the strain applied by the stretcher to the PDMS device. Our results show that even without a keratin network, the cells are well able to resist the equibiaxial strain.

### The keratin cortex protects the cells at intermediate strains

Based on previous studies that show stiffening of individual IFs at strains beyond 10 to 20%,^17^ the question arises of whether the mechanical properties of the WT and KO cells differ under strain. Hence, we perform force-mapping, i.e., combined AFM topography imaging with force spectroscopy measurements of the cell monolayers on PDMS membranes. Our stretcher is designed to fit on the stage of an AFM (Nanowizard 4, Bruker Nano GmbH). This setup enables simultaneous inverse light microscopy and AFM measurements (Fig. 2(a)). Figure 2(b, left) shows a typical topography image of a confluent cell monolayer, where the colors display the height differences within the sample from the lowest height (0 *µ*m, dark blue) to the highest point (here: 2.68 *µ*m, white). Note that this height difference is measured on top of the cell layer and not from the bottom of the cells. The image is post-processed by subtracting a plane to correct for tilt and additionally correcting each scan line individually by a subtracted slope.

**Figure 2:**
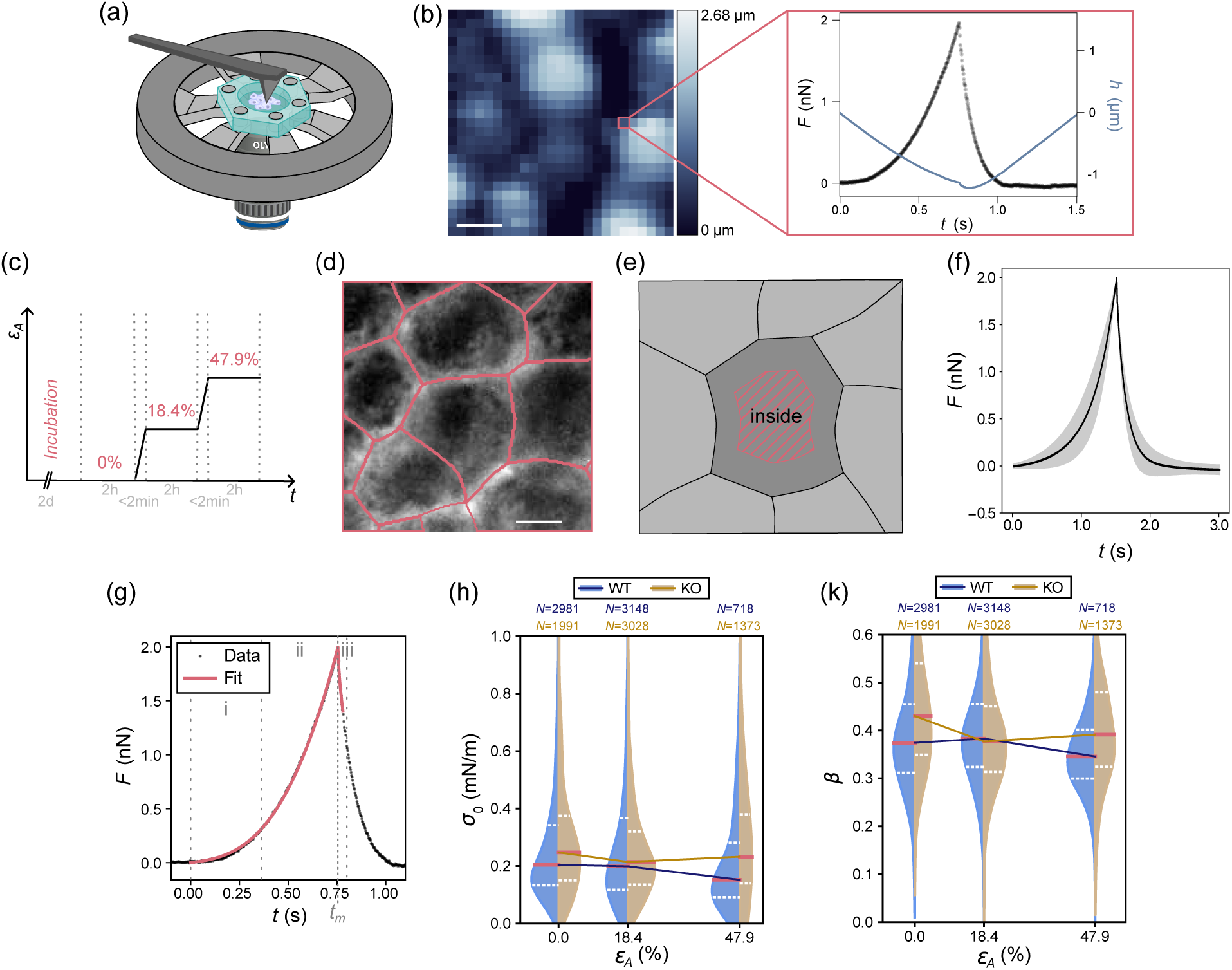
AFM indentation experiments. (a) Schematic of the cell stretcher with the AFM cantilever (dark gray) approaching from above, and the light microscopy objective from below. The cells (purple) are grown on the stretchable PDMS membrane (cyan). (b) (Left) Typical topography image of a confluent cell monolayer showing the sample height differences. Scale bar corresponds to 10 *µ*m. (Right) At each pixel, the cells are indented by changing the piezo height (blue) and a curve of the force *F* against time *t* is recorded (black). (c) Sketch of the timeline of a stretching experiment. The cells are seeded two days before the experiment. On the day of the experiment, each strain position is held for two hours, allowing us to image and indent the cells to perform AFM force spectroscopy. Reaching the next strain position takes less than two minutes. (d) Example phase contrast image of the cells with the segmentation (pink). Scale bar corresponds to 10 *µ*m. (e) Schematic of the inside region (pink hatched area) of the cell considered for the analysis. (f) Mean force curve (black, N = 736 individual data curves) and the standard deviation (gray shaded area) for the example of WT cells at 47.9% area strain. (g) An example plot of the force *F* against time *t* (black circles) together with a fit describing a viscoelastic model of the mechanical properties (pink line) and the different regions of the fit: low indentation depth (i), high indentation depth (ii) and relaxation curve (iii). The time of maximum indentation is marked with *t_m_*. (h) The pre-stress *σ*_0_ and (k) the fluidity *β* of MDCK II cells at different strains *ε_A_*. Data for WT cells are shown in blue, data for KO cells in yellow. The lines on the violins represent the median (solid pink line) and the upper and lower quartiles (dashed lines). The medians are connected via solid lines and the number of force curves *N* is displayed in the figure in the respective colors.

At each pixel, we perform a force spectroscopy measurement and a typical plot of the force *F* against time *t*, i.e., approach and retraction, is shown in Fig. 2(b, right, black). We indent the sample by lowering the piezo height of the AFM (Fig. 2(b, right, blue) up to a predefined force, the so-called set point. This maximum force is chosen such that the deformation of the cortex is not impaired by the nucleus or substrate. Hence, we ensure not to indent the sample more than 20% of the full sample height.^50^ For an average height of 10 *µ*m for MDCK II cells, this percentage corresponds to a value of 1-2 *µ*m.^2^ In comparison to typical indentation depths used for investigating the actin cortex,^50^ we indent slightly deeper to additionally also measure the keratin cortex and choose a set point of 2 nN. We use a spherical cantilever with 2 *µ*m diameter, which enables us to measure the mechanics of the cortex^50,51^ while keeping a high enough resolution for the imaging. Consequently, we choose a pixel size of the size of the cantilever, i.e., 2×2 *µ*m^2^. Of note, we manipulate the cells by applying a lateral strain with the cell stretcher (area change Δ*A* = *A*_new_ − *A*_0_), and then probe the viscoelastic properties of the cells by indenting with the AFM cantilever (area change Δ*S* = *S*_new_ − *S*_0_).

Figure 2(c) describes the timeline for a typical experiment. At each strain position, we take four force maps at adjacent positions with a size of 25 pixels × 25 pixels each, at a pixel size of 2×2 *µ*m^2^, resulting in one large image with a size of 100×100 *µ*m^2^. With an indentation velocity *v*_0_ of 2 *µ*m/s, each indentation measurement at every pixel takes approximately 3 s, summing up to a total time of around 2 h per strain position including any overhead time during change in strains. Increasing the strain takes less than 2 min. In total, a typical experiment takes 6 hours from starting the stretching experiment to finish when reaching the highest strain. Experiments are stopped prematurely, if the PDMS membrane breaks.

In addition to the force mapping, we take phase contrast light microscopy images of the cells at each strain position and segment the cells (Fig. 2(d)). By combining the AFM images and the light microscopy images, we can use the latter to segment the cells in the AFM images. To do so, we interpolate on the segmented outlines to fit the AFM pixel size. We overlay the outlines on the AFM image and manually translate it to align the cell outlines with the respective cells on the AFM image. Afterwards, we determine the “inside” region of the cells (Fig. 2(e)), which includes only the cell center that is up to three pixels away from the cell outline (SI Fig. S4). Altogether, we compare 6 conditions: two cell lines (WT, KO) and three strains (no strain, intermediate strain, high strain). Per condition, we collect 700 to 4000 force curves from 10 and 11 independent experiments for KO and WT cells, respectively.

We use the rating function implemented in the software package “nanite” ^44^ to exclude any failed curves from data acquisition. Additionally, we exclude those curves where the viscoelastic fit provides values that do not make sense from a theoretical point of view, such as values of the fluidity *β* smaller than 0, or an apparent compressibility modulus or pre-stress multiple orders of magnitudes larger than the expected values, that indicate a failed fitting of the curve. As expected for a complex biological system, we observe cell-to-cell variations as well as a distribution of the data curves even within one cell. All force curves are pre-

processed by correcting the baseline and performing a tip-sample correction to separate the distance of the cantilever moved towards the sample from the deflection of the cantilever in the opposite direction, aligned on the maximum, and averaged. Figure 2(f) displays a typical example of the mean force curve of all force curves within one condition (black) together with the standard deviation (shaded area). The force curves and their resulting average of all data categories are shown in SI Fig. S5 and S6.

To quantify the indentation data, we fit the force-time curves obtained from the AFM experiment using the viscoelastic Evans model, which is explained in detail in Refs.^52,53^ for a spherical indenter. This model takes into account the geometrical shape of the sample. For a cell in a monolayer, this shape is approximated as a spherical cap in the center of the cell, here called the “inside” region.

In the Evans model, the sample is defined via a generic shape function *g̃*_cap_ of a deformed sphere as a function of indentation depth. For a spherical indenter with radius *R_p_* on a spherical cap with radius *R*_1_, this function becomes

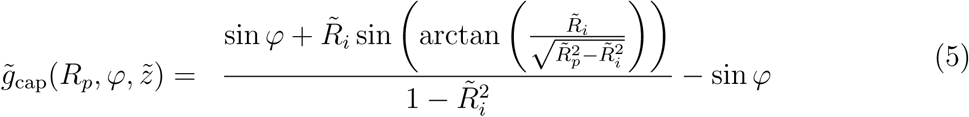

with *R̃_p_* = *R_p_/R*_1_ the dimensionless radius of the probe, *R̃_i_* = *R_i_/R*_1_ the dimensionless contact radius with the indenter and *φ* the contact angle changing with indentation depth *z̃* = *z/R*_1_. A sketch of the geometry can be found in the SI Fig. S7.

The cortex is assumed to resist deformations only by the tension *σ*. *σ* includes the pre-stress *σ*_0_, which the cortex naturally exhibits due to actomyosin contraction^6,54^ and the membrane attached to the cortex,^55,56^ as well as the area compressibility modulus *K_A_* as the response function to area dilation, assuming constant volume:^56,57^

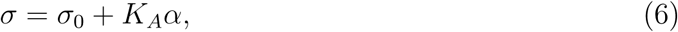

with the areal strain of the substrate 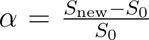, where *S*_0_ is the area before indentation and *S*_new_ is the area after indentation. The viscoelasticity is included through a power law for the compressibility modulus

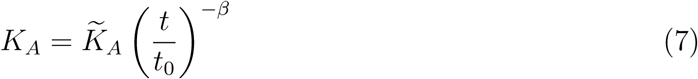

where *K̃_A_* is the apparent area compressibility modulus and *t*_0_ = 1 s (arbitrarily chosen).

The force response is then given by applying the elastic-viscoelastic-correspondence principle^28,52,58^ as

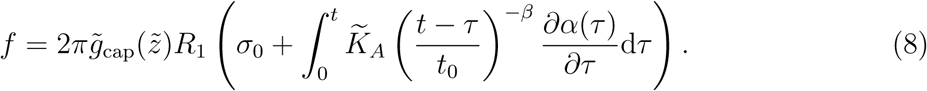

The integrals can be approximated through a polynomial 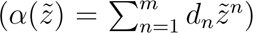 and divided into the force response to indentation, i.e., the approach curve *f*_app_(*t*) = *f* (*t* ≤ *t_m_*) until the time of maximum indentation *t_m_* (see Fig. 2(g)) and upon retraction *f*_ret_(*t*) = *f* (*t > t_m_*) starting at the time of maximum indentation *t_m_*. The resulting equations are used to fit the experimental force-time curves in a piece-wise manner (Fig. 2(g)) to obtain the pre-stress *σ*_0_, the scaling parameter or apparent area compressibility modulus *K_A_*, and the power-law exponent or fluidity *β*. 30% of the retraction part of the curve are included in the fit.

The pre-stress of the cortex is measured as the immediate response of the cell to the indentation at small times and indentation depths (Fig. 2(gi)), while the area compressibility modulus is measured when compressing the cell (Fig. 2(gii)). Finally, the fluidity is mainly determined through the relaxation part of the force response of the cell (Fig. 2(giii)), although the parameters are not independent of each other (Eqs. 6, 7).^53^

Figure 2(h) and (k) show the results for the pre-stress and fluidity at increasing area strains *ε_A_* as violin plots. The WT cells are displayed in blue, the KO cells in yellow. The solid pink lines on the violins represent the median of each distribution and the dashed lines the quartiles. As guide to the eye, the medians are connected with solid lines in the respective colors. The number of force curves is displayed in the respective color at the top of the figure. We find the pre-stress to be higher in KO cells, compared to the WT cells (Fig. 2(h)). This result first seems counter-intuitive as one may expect the pre-stress to decrease when the keratin is removed in the KO cells as there is less material that is deformed when indenting. However, as we use stable KO cells, it is likely that the cells react to the loss of keratin, for example, by alterations to the actin cortex and hence a change in the cortex contractility. Increased cortex contractility results in smaller cells as discussed in Ref. ^56^ and shown in Fig. 1(d), left, supporting this argument. Alternatively, but not contradictory, the cells might overcompensate the absence of the keratin with an increased amount of actin in the cortex, hence increasing the cortical tension. When comparing the pre-stress for the different strains applied to the cells, we find a decrease of *σ*_0_ from the relaxed state (0 % area strain) to the intermediate strain (18.4 % area strain) for the KO cells, whereas this decrease occurs only for the high strain (47.9 % area strain) in WT cells (Fig. 2(h)).

Likewise, the fluidity in WT cells changes little for intermediate strain and decreases at high strain, while in KO cells it strongly decreases already at intermediate strain and then remains at a similar value for the highest strain (Fig. 2(k)). Thus, the KO cells respond already to intermediate strains, whereas in WT cells the interplay between the actin and keratin networks preserve the original prestress and fluidity at these intermediate strains. The resilience of the actin network to the loss of keratin is remarkable, but at the same time, these results strongly indicate that KO cells mechanically compensate for the loss of keratin. Mechanical integrity, even under external influences is crucial for a cell to maintain its functions. For example, it has been shown that the cell can vary its surface area to keep the fluidity value constant.^28^ Additionally, keratin has been revealed to be the main component for mechanical resilience in keratinocytes.^11^ An increase in fluidity is often found in cancer cells or during wound healing.^59,60^ For the cell, it is important to remain deformable enough to adapt to changes but if the fluidity is too high the cell might lose the ability to keep its shape and functions. The actin cortex is well able to compensate for the loss of keratin and partially mimic the mechanical behavior of the WT cell. However, our findings indicate that, also in MDCK II cells, the keratin network plays an important role in ensuring cell integrity and preserving the overall mechanical properties of the cell and the keratin network stabilizes the cells by delaying the effect of stretching on the cells.

### 3D tissues from KO cells are more responsive to strain than WT tissues

As the KO cells exhibit a smaller cell size and increased pre-stress, as well as a more sensitive response to strain, we hypothesize that the actin cortex in the KO cells compensates for the loss of keratin. To investigate this hypothesis, we perform high-resolution imaging using STED microscopy of the actin cortices in 3D WT and KO tissues, see Fig. 3(b) and (c). MDCK II cells readily form 3D tissues when providing an extracellular matrix (ECM), such as hydrogels.^61^ In particular, the cells grow into so-called cysts by forming a spherical shell that encloses a lumen in the center. In a cyst, the apical side of the cell faces the lumen^62^ (see Fig. 3(a), cyan), in contrast to monolayers where the apical side forms the top of the cell layer. Because of the cyst’s spherical geometry, a lateral cross-section through its axial center reveals side views of the cells. In this orientation, the apical-basal axis lies within the imaging plane, enabling high-resolution imaging of the actin cortex along this axis with a standard STED microscope that improves lateral (but not axial) resolution. We first acquire a confocal microscopy image of the respective cross-section of the entire cyst (see Fig. 3(bi), (ci)) and then select the apical side of a single cell for high-resolution imaging (see Fig. 3(bii) and (cii)). We find that in the WT cells, the cortex is a thin homogeneously distributed structure, whereas the KO cells exhibit a more irregular structure with bundles of actin pointing outwards the cell towards the lumen (Fig. 3(cii), pink arrows). Hence, the KO cells indeed react to the loss of the keratin with a change in their cortical structure. This change might result in an increase in the actomyosin contractility, and consequently the pre-stress. These interesting results lead to the question of what the immediate response of the cell layers at the beginning of the stretching process is. With the equibiaxial stretcher, this process is not accessible, as it is necessary to realign the field of view during the stretching process to image and measure the mechanical properties of the same cells at varying strains, which leads to a delay in the experimental process. Additionally, as we grow the cells into a confluent monolayer in the PDMS device, we only investigate a small section of the tissue. To overcome these limitations, we develop a complementary method to stretch whole MDCK II cells, grown into cysts, fast to high equibiaxial strains while imaging the entire tissue throughout the full process.

**Figure 3:**
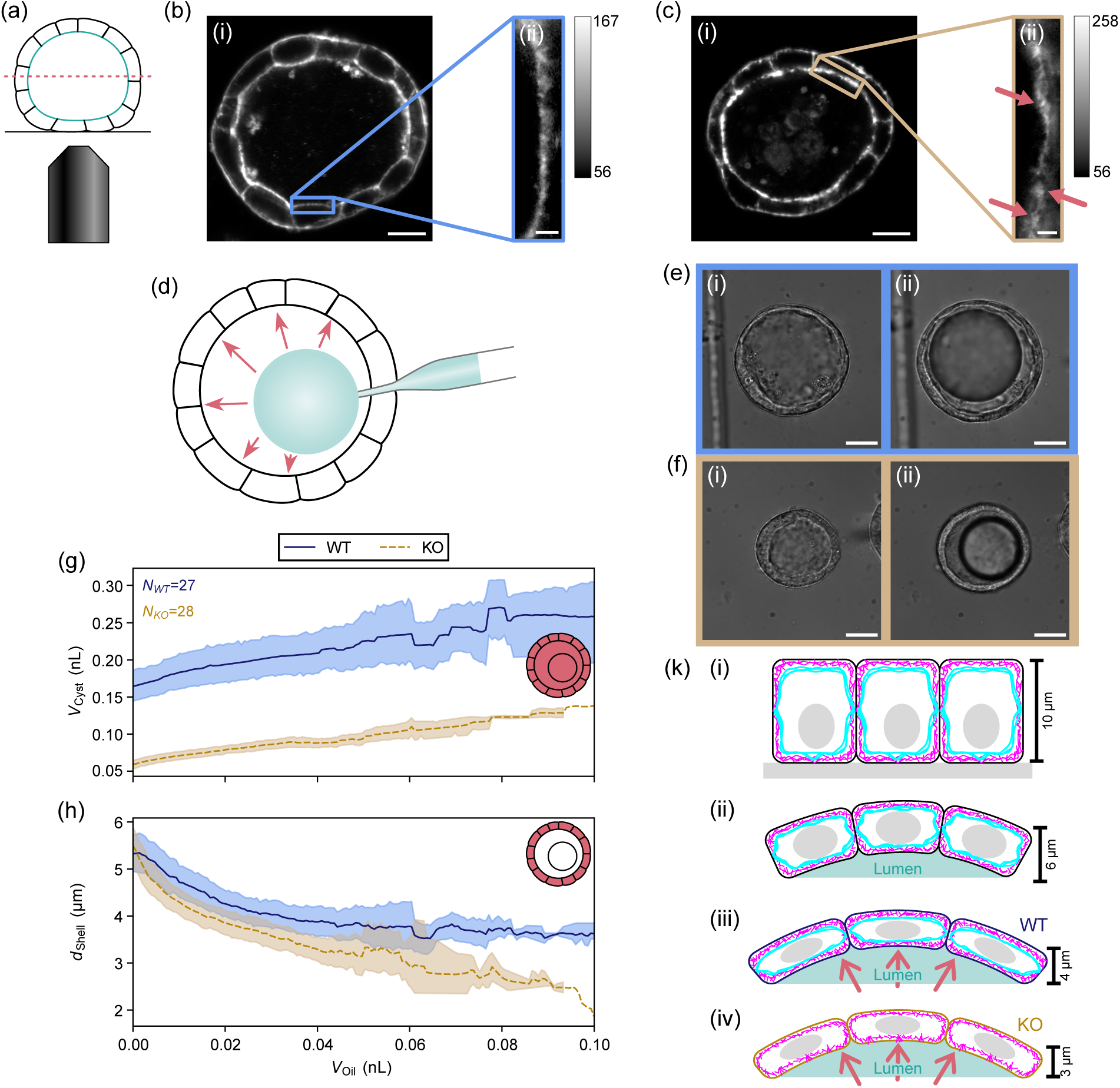
3D cysts grown from MDCK II cells. (a) Schematic side view of a cyst and the imaging plane (dashed pink line). The apical side of the cells (cyan) is facing the lumen. (b-c) Fluorescence images of the actin cortex in MDCK II cysts. (b) WT cyst. (c) KO cyst. (i) Confocal microscopy images of the lateral cross-section of a cyst. Scale bars correspond to 10 *µ*m. (ii) STED microscopy images of the apical actin structure in a single cell. The pink arrows point towards irregular structures of the cortex in the KO cyst. Scale bars correspond to 1 *µ*m. The gray scale bar shows the recorded signal in arbitrary units. (d) Schematic of the cyst injection. The micropipette (gray) is inserted into the cyst and the oil (cyan) is released into the lumen of the cyst (black) hence stretching the cyst (pink arrows). (e-f) Example phase contrast microscopy images of the WT and KO cells as cysts. (e) Living WT cyst. (f) Living KO cyst. (i) Unstrained cyst. (ii) Strained cyst. Scale bars correspond to 20 *µ*m. (g) Cyst volume *V*_Cyst_ (pink area in icon) plotted against increasing oil volume *V*_Oil_. (h) Shell thickness *d*_Shell_ (pink area in icon) plotted against increasing oil volume *V*_Oil_. Results for WT cells are displayed in blue with a solid line, for KO cells in yellow with a dashed line. Lines are showing the mean of all data sets. The shaded area represents the 95% confidence interval. The number of cysts *N* is displayed in the figure in the respective color. (k) Comparison of the cell thickness in each regarded case, drawn to scale as obtained from the experiments: (i) in a monolayer, (ii) cyst and (iii) stretched WT and (iv) KO cysts. The actin and keratin networks are shown in magenta and cyan, respectively.

We increase the intraluminal pressure of living cysts using microinjection to enforce a stretching of the cell shell. Inspired by Wang, Zhao et al.,^63^ we inject insoluble mineral oil into the lumen of the cysts to stretch the cells (see Fig. 3(d)). By stretching the cysts through oil injection, we apply the strain on the cells from the apical side. As it was shown before that the apical side of the cells is softer and dissipates the energy through the tissue to prevent damage, while the basal side of the cells provides elasticity to the system,^64^ the stretching from the apical side might result in a different response of the cells than monolayer stretching. Furthermore, we image the side-view of the full tissue that is exposed to strain in comparison to the small field of view from the top when imaging the monolayer. Therefore, the combination of both stretching methods and imaging approaches offers complementary information.

We seed the cells 7–9 days in advance in Matrigel to allow for cyst formation. Right before the experiments, we remove the Matrigel from the cysts using “cell recovery solution”. We perform the injection on an inverse microscope which allows us to take phase contrast videos during the complete injection process. To stretch the cyst, we carefully insert a micropipette (see Fig. 3(d), gray) into the cyst. We manually control the amount, rate and pressure of mineral oil (cyan) that is released into the lumen (pink arrows). The injection process typically takes 1-3 s. Finally, the pipette is pulled out of the cyst. To investigate the influence of keratin on the response of the cells to stretching, we use the same WT and KO cell lines are before. Figure 3(e) and (f) show example phase contrast microscopy images of WT and KO cysts, respectively: unstrained, relaxed cyst (i), strained, injected cyst (ii). Example videos of the injection of a WT and a KO cyst are provided as SI Movie 1 and 2. We perform 10 independent experiments each, where one Petri dish with multiple cysts is considered one experiment. We inject oil into 27 WT and 28 KO cysts, respectively.

A visual inspection of the phase contrast images of both WT and KO cysts reveals that the KO cysts are smaller than the WT cysts. To rule out effects of the growth time on the size of the cysts we compare both cell lines at the same time span range after seeding. A manual estimation conducted by counting the nuclei in the imaged plane (Fig. 3(a)) confirms that the cysts in both cell lines contain approximately the same number of cells.

To investigate the response of WT and KO cysts to stretching, we determine the cyst shape at increasing oil volumes. The analysis workflow is schematically shown in SI Fig. S8. We manually segment the cyst, lumen and oil droplet individually in each single frame of the video from one frame before the first oil droplet is released into the lumen until the size of the oil droplet does not change any further. We determine the center of mass of the lumen and calculate the distance from the center of mass to the outline of the lumen and the cyst individually for varying angles. By averaging over all distances, we obtain the average radii *r* for the lumen and cyst. We repeat the procedure for the oil droplet separately. In first approximation, we assume the cyst and the oil droplet to be spherical and determine their volume *V* . The shell of the cyst is defined as the cyst without the lumen, hence the thickness of the shell *d*_Shell_ is given by Eq. 2. As the injection pressure and time are manually controlled, and therefore arbitrary, we compare all cysts by the injected oil volume.

Figures 3(g) and (h) show the cyst volume *V*_Cyst_ and shell thickness *d*_Shell_ with increasing oil volume *V*_Oil_. The WT cells are displayed in blue by a solid line and the KO cells in yellow by a dashed line. The lines display the mean over all cysts and the 95% confidence bands are presented as the shaded areas in the respective colors. SI Figure S9 shows the same data normalized by the volume of the cyst *V*_Cyst,0_ and the shell thickness *d*_Shell,0_ before injection to compare relative changes.

In agreement with our visual impression, the quantitative analysis reveals that the KO cysts are indeed smaller than the WT cysts (see Fig. 3(g)). This result supports our results from above that the actin cortex in the KO cells exhibits a higher contractility as an increase in the tension also increases the pressure on the lumen and consequently resulting in a smaller lumen, see Fig. 3(e), (f). Interestingly, however, the thickness of the shell is the same in relaxed KO cysts and WT cysts, see Fig. 3(h). Hence, keratin KO cells are well able to form a spherical lumen with a normal shell. Both WT and KO cysts increase in size with the injected oil volume and the shell becomes thinner, indicating a stretching of the cells, assuming a constant volume. This volume change translates to a surface area increase of the cyst of up to 32%, indeed overcoming the limit of the cell stretcher with the maximum cell area increase in our experiments of 23%. When comparing the change in size with increasing strain applied by the oil droplet, we find that less oil is needed to stretch the KO cysts by the same relative amount than in the WT cysts, see SI Fig. S9(a).

It has been shown in in vitro experiments, that actin ruptures at relatively low forces, while IFs can resist high strains^14,15,18^ and in stretched cells, the actin network fluidizes. ^26,28^ Hence, the actin network alone is not able to keep the cellular tension at a physiological level, which results in highly compliant cells. In the WT cysts, however, the keratin network can withstand those forces due to its extensibility and preserves the cyst from rupturing at high strains. It was found that in cell domes the keratin network reorganizes into bundles that allow to “superstretch” the cell. When grown into domes, the apical side of the cells is facing upwards, as in the monolayer. These keratin bundles were also shown to be load bearing: cutting the keratin bundles results in an immediate increase in cell size.^37^ Moreover, IFs are often referred to as the safety belt of the cells as they are easily deformable at slow deformations but stiffen upon fast deformations. ^17,22–24^ In contrast to Latorre et al.^37^ where the dome forms within multiple hours, we stretch the cysts on the timescale of seconds, thus addressing the regime of “fast deformations”. Hence, the keratin network presumably stiffens when the cell is stretched to protect the cell and preserve its integrity.

We observe that the shell thickness of the KO cysts decreases more strongly than in the WT cysts, see Fig. 3(h). With a shell thickness of around 5.5 *µ*m, the cysts form thinner cell sheets approximately half the thickness than a monolayer with around 10 *µ*m height for MDCK II cells,^2^ see Fig. 3(ki) and (kii), possibly due to the intraluminal pressure of cysts.^37^ When increasing this pressure through the injection process, the cells get stretched and hence become thinner. Interestingly, the shell thickness reaches a plateau in the WT cysts at *d*_Shell_ ≈ (3.83 ± 0.06) *µ*m (±SE), while the KO shells continue to thin, see Fig. 3(h) and 3(kiii) and (kiv), respectively. However, it has to be noted that there are too few curves at large oil volumes to be statistically relevant. In both cell lines, other cellular components, such as the nucleus, mechanically limit the thinning of the cell. Any further thinning would lead to a rupture of cell. From these results we hypothesize that in the WT cells, where the keratin network forms the circumferential rim together with the radial spokes and a cage around the nucleus,^9^ this keratin network might restrict the thinning process to the plateau value to protect the cell and nucleus from rupture. In the KO cells, this perinuclear keratin cage is missing leading to a further thinning of the cells.

We show that the WT cysts are more resilient to stretching than the KO cysts. The keratin network in the WT cells preserves the cell integrity and protects the cellular components from damage. Without a functioning keratin network, the cells cannot cope with the forces when stretched to high strains which leads to a more pronounced stretching and thinning of the cells and consequently to a rupture when stretched further. Our results suggest that the cells are well able to survive without a keratin network and form into 2D and 3D tissue, but at high strains the extensibility and resilience of the keratin network is needed to protect the cell from damage.

### The keratin cortex ensures cell integrity at high strains

While these results demonstrate the overall importance of the keratin network for cellular integrity under strain, they do not address whether its protective effect varies within different cellular regions. As the MDCK II cells express keratin in a rim-and-spokes-like arrangement,^9^ the keratin network might add additional mechanical properties to the cell when comparing the periphery of the cells to the cell center. Hence, from the segmentation of the cells in our AFM data, we include an additional region and we divide each cell now into two regions: “rim” and “inside” (Fig. 4(a)). From the segmentation, we obtain the outline of the cells right at the outermost pixels that are still included in that cell. The rim is defined as exactly this one pixel wide line. The inside region is defined as before, the cell center up to three pixels away from the rim. In-between both regions, we exclude an interim area between the two regions to avoid any unclear classifications (SI Fig. S4).

**Figure 4:**
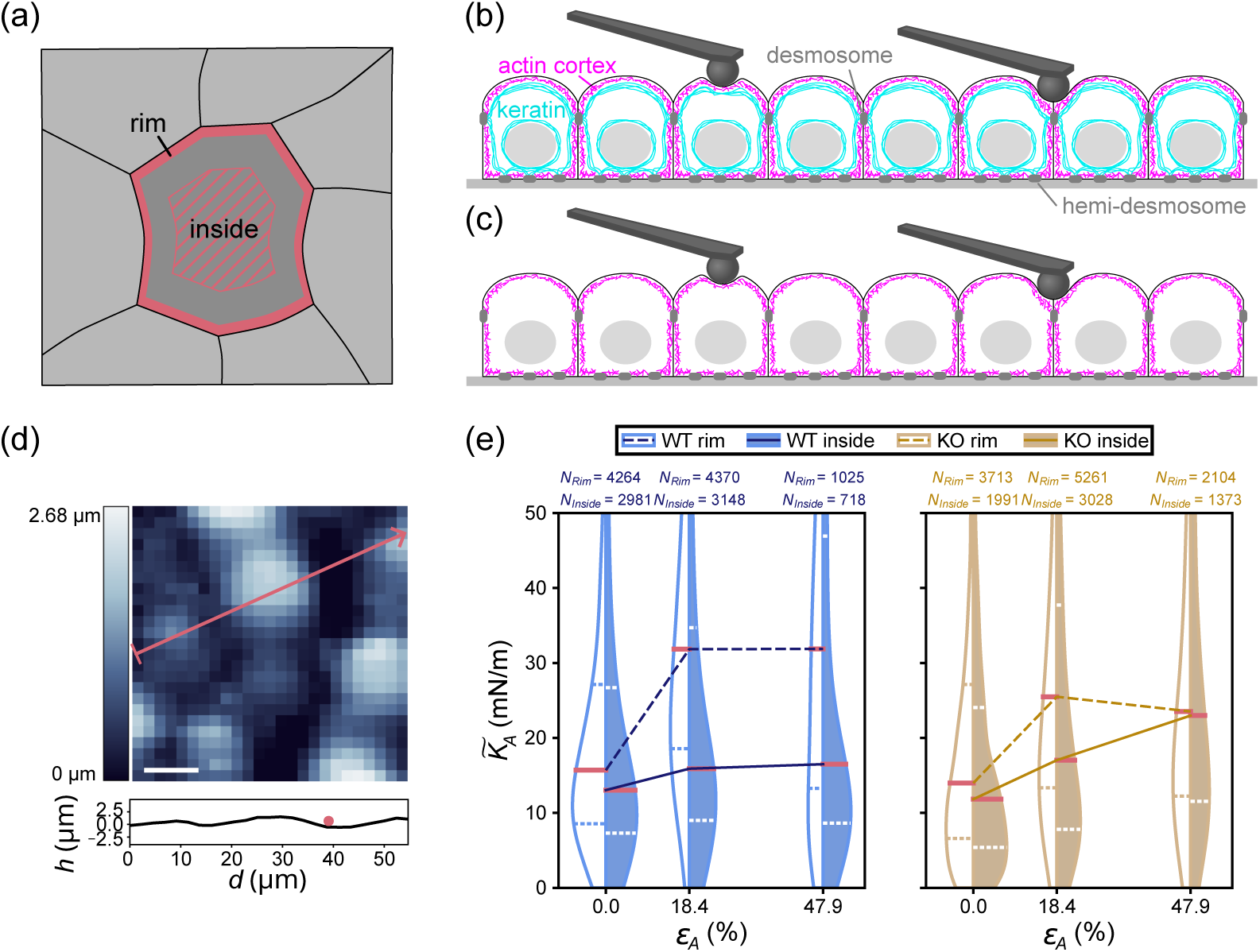
Viscoelastic response of different cell regions. (a) Schematic of the division of the cell area into two regions: rim (thick pink line) and inside (pink hatched area). In-between both regions, an interim section is excluded from the analysis. (b) WT, (c) KO cells; schematic side view (not to scale) of the actin cortex (magenta), keratin network (cyan), desmosomes and hemi-desmosomes (dark-gray) and nucleus (gray) in WT cells that are indented with a cantilever (black) in the cell center (left) and at the cell-cell contacts (“rim” of the cell) (right). (d) An example line profile (pink line) of the height of the cells is shown together with the spherical indenter of radius 1 *µ*m (pink circle) drawn to scale. Scale bar corresponds to 10 *µ*m. (e) The apparent compressibility modulus *K_A_* of MDCK II cells at different strains *ε_A_*. Data for WT cells are shown in blue, data for KO cells in yellow. Filled violins correspond to the inside-region of the cells, open violins with colored outlines to the rim-region. The lines on the violins represent the median (solid pink line) and the upper and lower quartiles (dashed lines). The medians are connected via lines: dashed lines for the rim-region and solid lines for the inside-region. The number of force curves *N* is displayed in the figure.

Fig. 4(b), (c) represent schematic side views (not to scale) of cells containing an actin (magenta) and a keratin (cyan) cortex (b) or only an actin cortex (c) and the indentation of the two different regions, i.e., the inside (left) and the rim (right) regions. An example 3D microscopy image of fixed WT cells shows this structure (SI Fig. S10). In the inside region, we indent the cortex that is oriented perpendicularly to the indenting direction, forming a thin sheet. In the rim region, the keratin cortex connects to the desmosomes and is oriented in parallel to the indentation direction. As a result, the cantilever effectively indents more keratin and compresses the network in vertical direction and we expect a difference between the regions mainly in the stiffness of the cells.

Although the cell shape is approximated as a spherical cap in the Evan’s model, for a cell curvature that is large compared to the size of the cantilever, the cantilever encounters a close-to-flat surface. Hence, we apply this model also to the rim region of the cell, because the diameter of the spherical indenter (2 *µ*m) is small compared to the curvatures on top of the cell layer (see Fig. 4(d) for a line profile of the height of the cells and a cantilever (pink closed circle) schematically drawn to scale).

Figure 4(e) shows the apparent area compressibility modulus at increasing area strains *ε_A_* as violin plots. The compressibility modulus describes how much pressure is needed to achieve the compression of the cell and is hence a measure for the stiffness of the cell. The WT cells are displayed in blue, the KO cells in yellow. Filled violins represent the results from the inside region, while open violins represent the results for the rim region. The solid pink lines on the violins represent the median of each distribution and the dashed lines the quartiles. As guide to the eye, the medians are connected with solid lines for the inside region and dashed lines for the rim region in the respective colors. The number of force curves is displayed in the respective color at the top of the figure. Further violin plots comparing different pairs of data sets and the pre-stress as well as the fluidity are shown in SI Figs. S11-S13 and dataset S1.

We find the area compressibility modulus *K_A_* to be increased in the rim region of both WT and KO cells compared to the inside region in the relaxed state (Fig. 4(e)). In the rim of the cells, more force is needed to compress the cortex as here the cantilever not only deforms the cortical networks that form a thin sheet, perpendicular to the indentation direction, but also compresses the part of the cortex that lays in parallel to the indentation direction, i.e., the adherens belt (Fig. 4(b), (c) right). Hence, there is more effective cortex in the rim region both in the cells with and without keratin increasing the stiffness on the cell-cell contacts. Remarkably, the difference between rim and inside is much more pronounced for the WT cells, which can be attributed to the stiffening of IF under compression as shown for vimentin in vitro and in cells.^25^

In the inside region, the cells are easier to deform, meaning that the cell is softer. This result supports our assumption above of less effective cortex compared to the rim region. Furthermore, the compressibility modulus of the KO cells increases steadily, while in the WT cells the modulus only increases from the relaxed state to the intermediate strain and is preserved from the intermediate strain to the high strain (Fig. 4(e)). Cells stiffen during various processes, such as cell division or movement presumably through an activation of the myosin motors or cross-linking of the networks.^65,66^ Both actin filaments and IFs are known to contribute to the stiffening.^65^ At intermediate strains, the cell stiffens to stabilize the cell, whereas at high strains, this stiffening does not continue further likely to keep the cell’s ability to deform and adjust to internal and external changes. It is highly remarkable that in the keratin KO cells the actin cortex is able to compensate for the loss of keratin to fulfill the same purpose of stiffening, although the stiffness is decreased in the rim region. However, at high strains, when the actin filaments start to rupture^14^ and the actin network fluidizes, the interplay of the keratin network and the actin cortex is necessary to avoid a further stiffening to preserve the functions of the cell.

## Conclusion

We present two complementary methods to investigate the effect of stretching on cells: an equibiaxial cell stretcher and microinjection of cysts. We study two lines of MDCK II cells, i.e., with (WT) and without (KO) keratin. We find that the keratin-deficient cells are well able to form tissues, both monolayers and cysts, and they follow and survive the high strains applied here, similar to the WT cells. Hence, the cells can compensate for the loss of keratin, however high-resolution imaging shows that the actin cortex structure in the KO cells is altered. Remarkably, the unstretched KO and WT cells are smaller but both exhibit the same thickness, as we see in the shell thickness of the cysts, hence the total volume of the KO cells is decreased.

The equibiaxial stretcher is compatible both with light and atomic force microscopy, enabling us to measure the mechanical properties of the sample while imaging the cells. We perform force mapping experiments, combining AFM imaging with spatially resolved force spectroscopy and overlay the AFM images with the light microscopy images to divide the cells into two regions according to the rim-and-spokes hypothesis. This approach enables us to compare the mechanical properties of the two regions. By fitting a dedicated model to the data, we determine the viscoelastic properties of WT and KO cells to investigate the influence of the keratin cortex to cell mechanics. Our results suggest that in KO cells the actin network compensates for the missing keratin network, leading to an increase in pre-stress compared to the WT cells. This result is supported by the fact that the KO cells are smaller and form smaller cysts.

We compare an “intermediate” and a “high” area strain. For both the pre-stress and the fluidity we observe that the KO cells display altered mechanical properties already at the intermediate strain, whereas for WT cells, the actin and keratin networks in concerted action are able to maintain the original, i.e., unstrained, values and react only to the high strain. Similarly, the KO cysts react more sensitively to the stretching process than the WT cysts, proving the necessity of the keratin network to stabilize the cells under strain. Our results highlight the importance of the keratin network and more specifically the keratin cortex for the integrity of the cell.

## Acknowledgement

We thank Ruben Haag, Peter Luley, Angela Rübeling, Mihaela Raycheva, Greta Höhndorf, Marie Tersteegen, Axel Munk, Timo Betz and Jonathan Frohn for technical support and fruitful discussions. This work was funded by the Deutsche Forschungsgemeinschaft (DFG, German Research Foundation): project-ID 449750155 - RTG 2756, projects A2, A7 (to S.K.), projects A6, B1 (to A.J.); project-ID 430255655 - KO3572-8/1 (to S.K.); project-ID 449544493 (to S.K.); project-ID 390729940 - EXC 2067/1 (to S.K. and A.J.). The work was further financially supported by the European Research Council (ERC, Grant No. CoG 724932, to S.K.) and was conducted within the Max Planck School Matter to Life supported by the German Federal Ministry of Education and Research in collaboration with the Max Planck Society.

## Author contributions

S.K. conceived and supervised the project; R.M. performed the AFM experiments; R.M. and U.U. performed cyst stretching experiments; RM. and P.J. performed STED imaging; S.K., A.J. and C.G. supervised the work; R.M. analyzed the data; U.R. prepared the fluorescently tagged cell lines and performed the cell culture; N.S. prepared the KO cell line; A.P.-T. provided analysis code; A.V.S. characterized the cell-stretcher; A.J. developed the viscoelastic model; R.M., S.K. and A.J. interpreted the results; R.M. and S.K. wrote a first draft of the manuscript, all authors contributed to writing the manuscript.

