## Supplementary Information for "The Keratin Cortex Stabilizes Cells at High Strains"

### Supplementary Movies

File Name: Movie_1

Description: Typical phase contrast video of the injection of the WT cyst presented in Fig. 3e in the main text. The scale bar corresponds to 20 $\mu$m, the time is displayed in the movie.

File Name: Movie_2

Description: Typical phase contrast video of the injection of the KO cyst presented in Fig. 3f in the main text. The scale bar corresponds to 20 $\mu$m, the time is displayed in the movie.

### Supplementary Dataset

File Name: Dataset_S1.csv

Description: The results from the viscoelastic fitting of the AFM force curves.

### Supplementary Figures

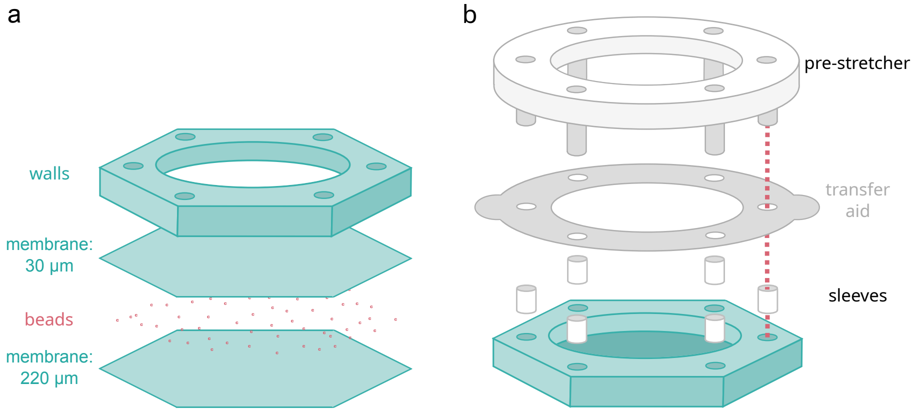


Figure S1: Design of the PDMS devices and pre-stretcher. (a) The hexagonal PDMS device consists of a membrane and wall attached to it. The membrane is made of two layers of PDMS with fluorescent beads in-between. The lower part of the membrane is approximately 220 $\mu$m thick, the top layer 30 $\mu$m. The PDMS walls form a well for the cells. Six holes are punched through the finished device on all six corners of the hexagon. (b) The pre-stretcher keeps the device flattened out during cell culture. The six holes in the devices are stabilized by PTFE sleeves. The pre-stretcher inserts into the sleeves with six pins. A transfer aid is inserted between the pre-stretcher and the device enabling easy transfer from the pre-stretcher on the stretcher for experiments.

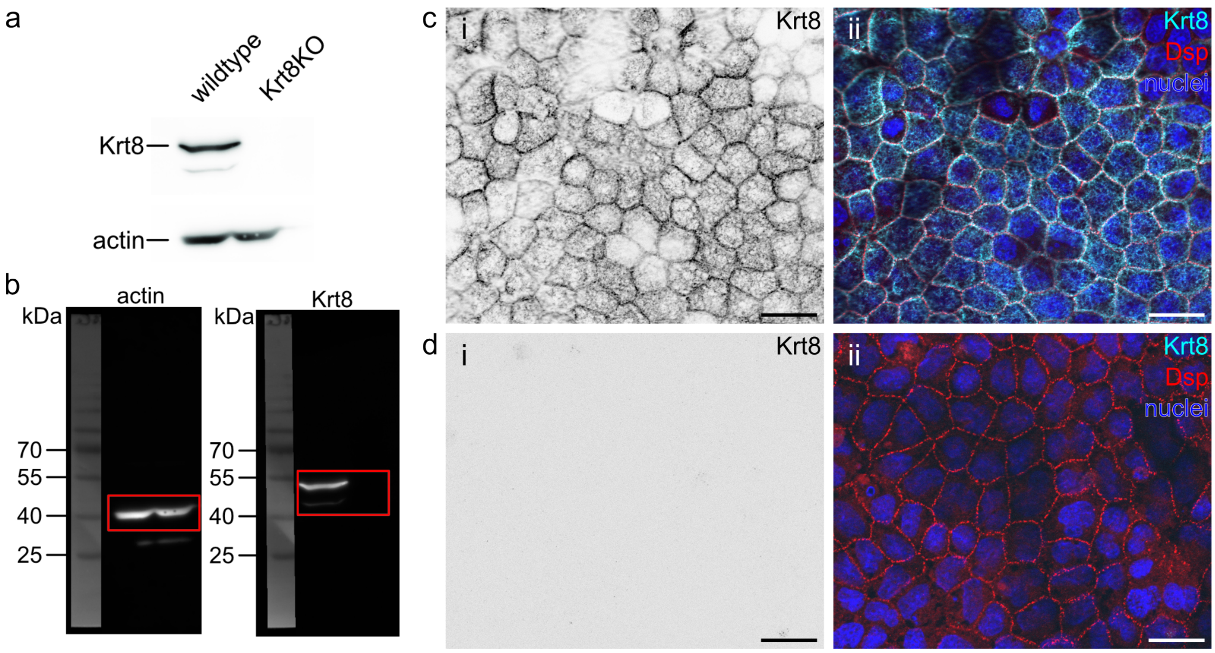


Figure S2: Confirmation of successful keratin 8 knock-out. (a) Immunoblot analysis of original wild-type MDCK II and keratin 8 knock-out (KO) cells. (b) Raw data from (a). (c-d) Staining of (c) wild-type and (d) KO cells. (i) Staining of keratin 8. (ii) Corresponding composite images showing nuclei (blue), desmoplakin (red) and keratin 8 (cyan) in wild-type and KO cells. Scale bars correspond to 20 $\mu$m.

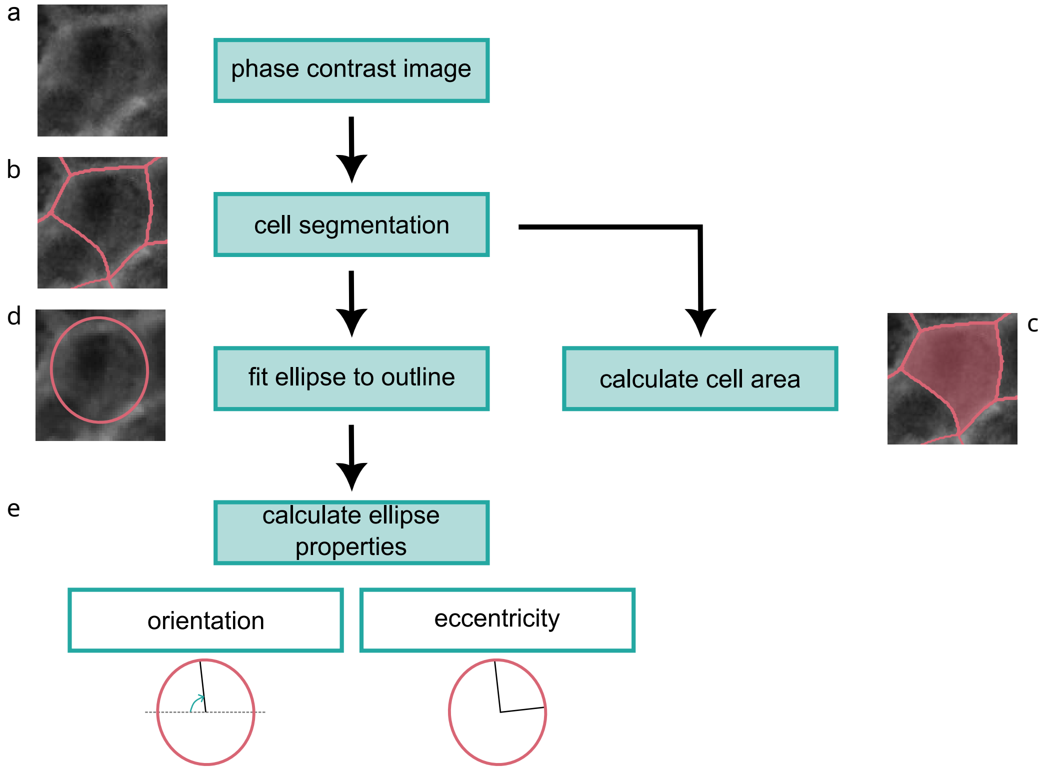


Figure S3: Workflow for the analysis of the cell shape. (a) The phase contrast image of the cells is (b) segmented to get the outline of each cell and (c) the area per cell is determined. (d) An ellipse is fitted to the outline and (e) the orientation and eccentricity are calculated.

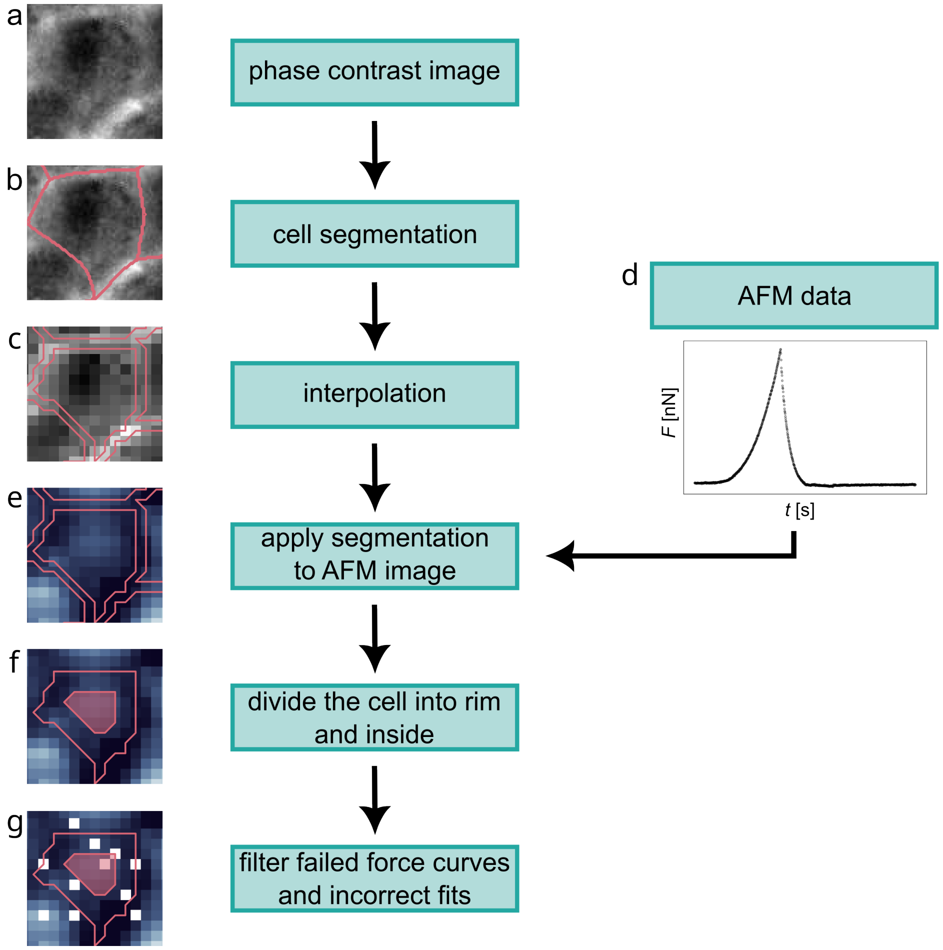


Figure S4: Workflow for the analysis of the force mapping data. (a) The phase contrast image of the cells is (b) segmented to obtain the outline of each cell. (c) The outline is interpolated to a bigger pixel size and (e) overlayed to the AFM image, (d) consisting of the force spectroscopy curves at each pixel. (f) The cell is divided into the rim and the inside regions and an interim section is excluded. (g) Failed curves and outliers are excluded.

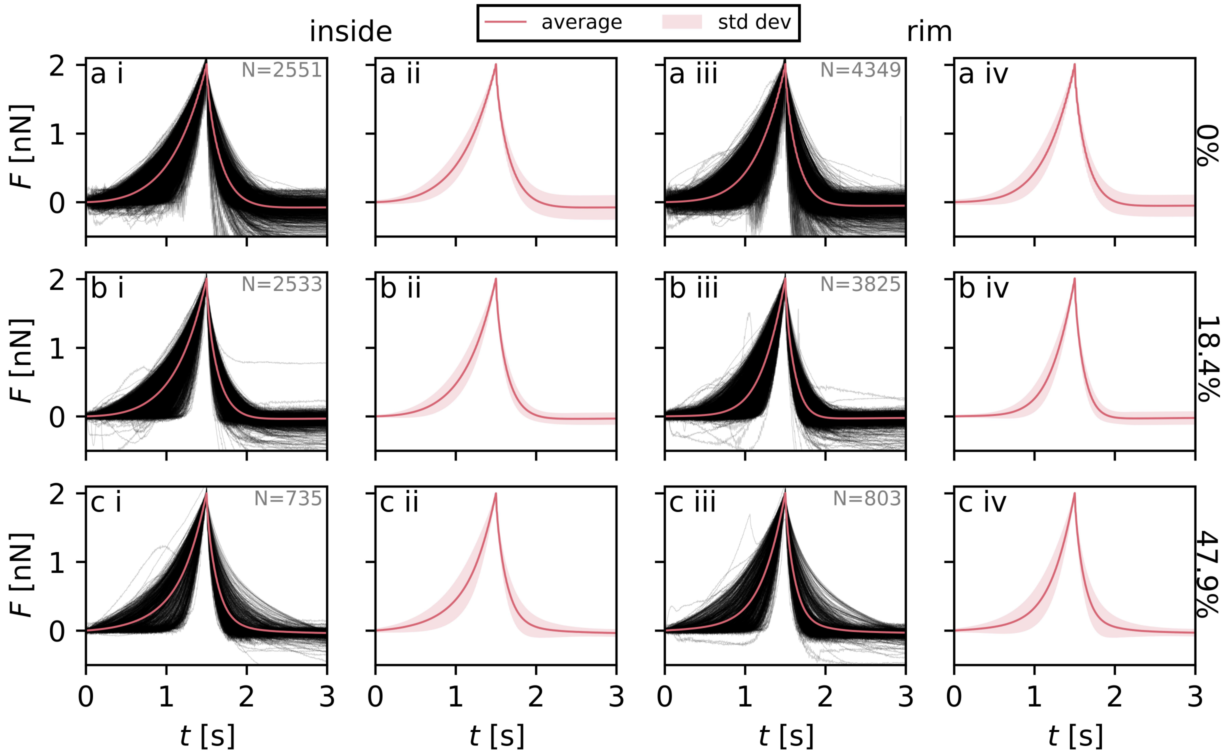


Figure S5: Force-time curves of the WT cells for an area strain of (a) 0%, (b) 18.4% and (c) 47.9%, for (i, ii) the inside region and (iii, iv) rim region. (i, iii) All single force curves (black) and their average (pink). (ii, iv) The same averaged force curve as in i and iii respectively (pink) and the standard deviation (pink shaded area). The number of force curves $N$ is provided in the figure.

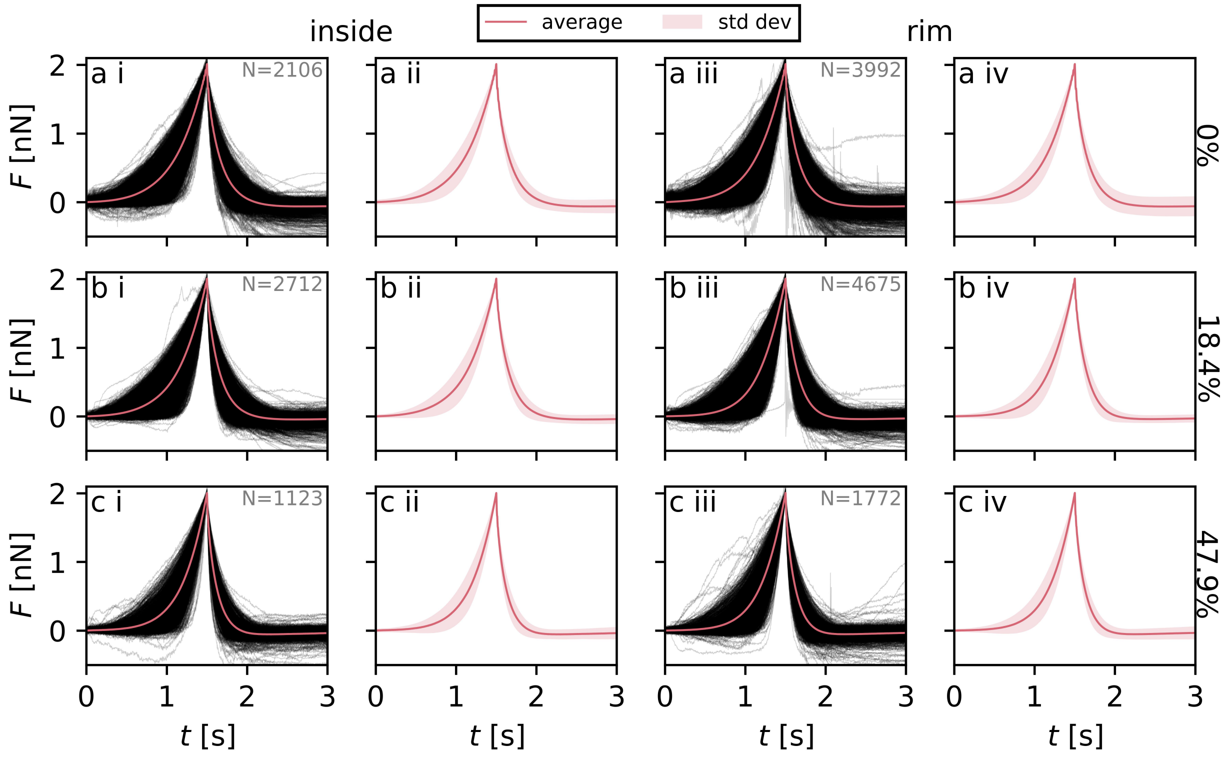


Figure S6: Force-time curves of the KO cells for an area strain of (a) 0%, (b) 18.4% and (c) 47.9%, for (i, ii) the inside region and (iii, iv) rim region. (i, iii) All single force curves (black) and their average (pink). (ii, iv) The same averaged force curve as in i and iii respectively (pink) and the standard deviation (pink shaded area). The number of force curves $N$ is provided in the figure.

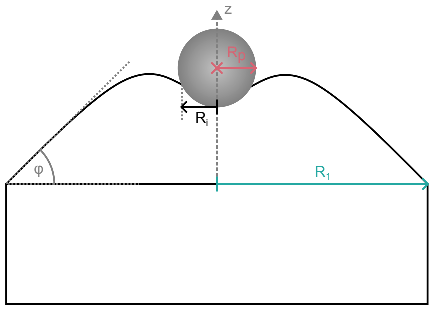


Figure S7: Schematic of the geometry for the Evans model. A cell in a monolayer is described as a spherical cap with radius $R_1$ and contact angle $\varphi$. During indentation with a sphere with radius $R_p$ by indentation depth $z$, the contact can be described by the contact radius $R_i$.

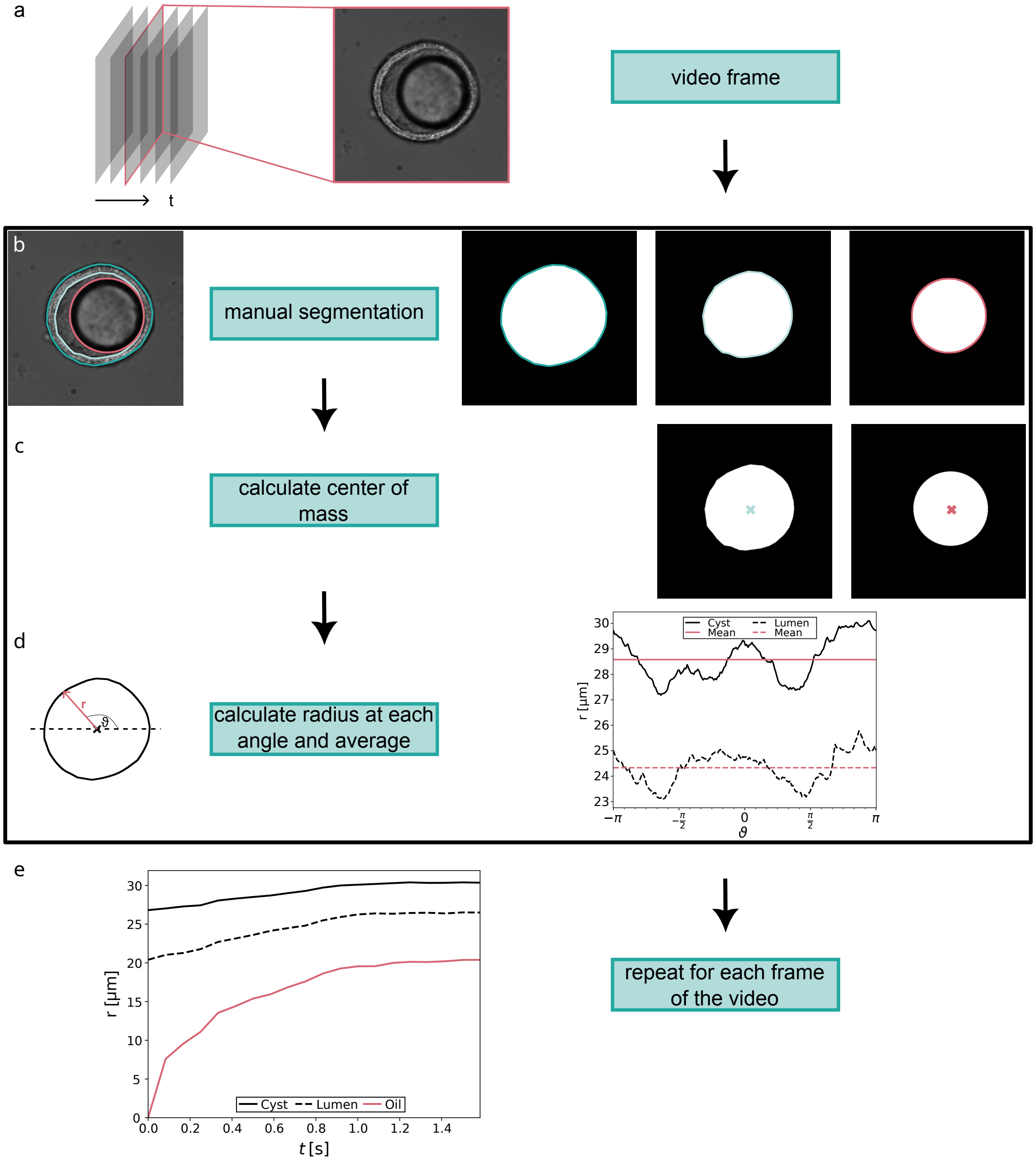


Figure S8: Workflow for the analysis of the cyst injection. (a) The video is split into individual frames from one frame before the first oil droplet is injected until the oil droplet does not change size anymore. (b) In each frame, the cyst (dark cyan), the lumen (light cyan) and the oil droplet (pink) is manually segmented and transformed into a mask. (c) The center of mass is calculated for the mask of the lumen and the oil droplet individually. For the cyst mask, the center of mass from the lumen is taken. (d) The radius from the center of mass to the outline of the mask is determined at various phase angles. The radii are individually averaged to generate one radius for the cyst, lumen and oil droplet respectively. (e) The procedure is repeated for each frame of the video.

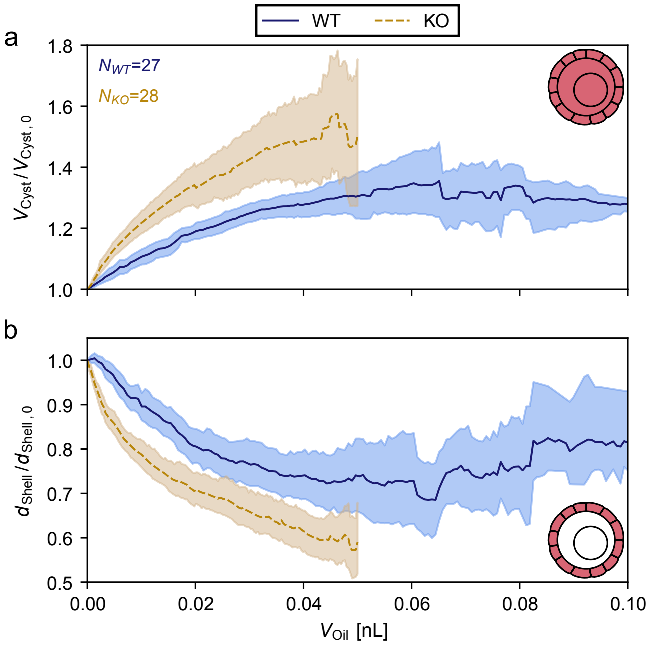


Figure S9: The normalized volume and shell thickness of the cysts upon stretching. (a) Normalized cyst volume $V_{\mathrm{Cyst}}/V_{\mathrm{Cyst},0}$ (pink area in icon) plotted against increasing oil volume $V_{\mathrm{Oil}}$. (b) Normalized shell thickness $d_{\mathrm{Shell}}/d_{\mathrm{Shell},0}$ (pink area in icon) plotted against increasing oil volume $V_{\mathrm{Oil}}$. Results for WT cells are displayed in blue with a solid line, for KO cells in yellow with a dashed line. Lines are showing the mean of all data sets. The shaded area represents the 95% confidence interval. The number of cysts $N$ is displayed in the figure in the respective color. For the KO cells, we cut the data at an oil volume of 0.05 nL, as we have too few curves for higher volumes which results in high noise when normalized.

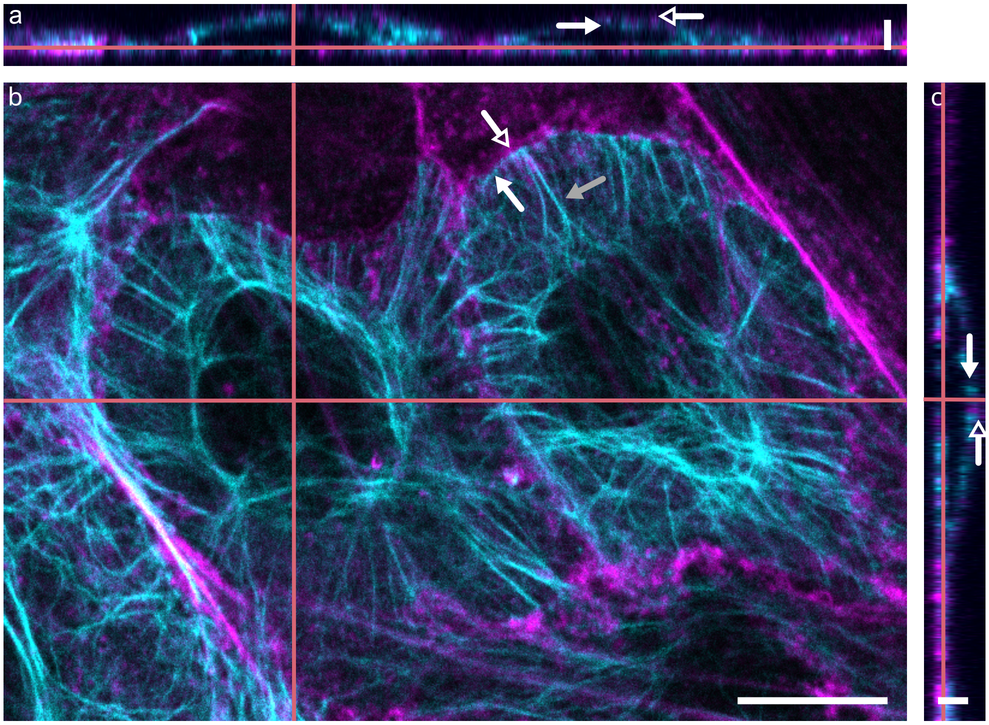


Figure S10: Example 3D confocal microscopy image of keratin tagged with EGFP (cyan) and lifeact actin tagged with mCherry (magenta) in fixed MDCK II WT cells. Pink lines indicate the displayed $x$, $y$, and $z$ slices. (a) $xz$-view. Scale bar shows the $z$-scale and corresponds to $2\,\mu$m. The keratin cortex (closed arrow head) lies underneath the actin cortex (open arrow head). (b) $xy$-view. Actin cortex (open arrow head) and keratin rim (closed arrow head, white) and spokes (closed arrow head, grey). Scale bar shows the $x$- and $y$-scale for all three images and corresponds to $10\,\mu$m. (c) $yz$-view. Scale bar shows the $z$-scale and corresponds to $2\,\mu$m. The keratin cortex (closed arrow head) lies underneath the actin cortex (open arrow head). Note that the cells appear particularly flat, because they are chemically fixed and prepared between two glass slides.

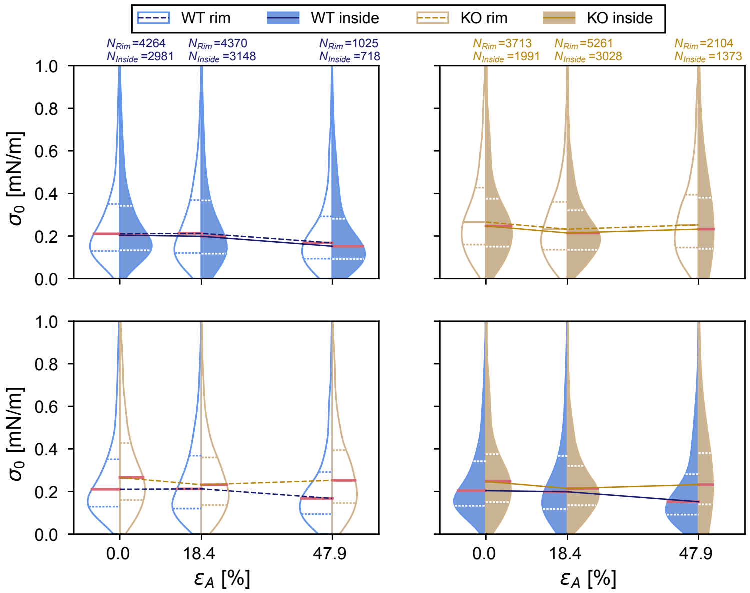


Figure S11: Pre-stress $\sigma_0$ of cells under strain $\varepsilon_A$. Data for WT cells are shown in blue, data for KO cells in yellow. Filled violins correspond to the inside-region of the cells, open violins with colored outlines to the rim-region. The lines on the violins represent the median (solid pink line), the upper and lower quartiles (dashed lines). The medians are connected via lines: dashed lines for the rim-region and solid lines for the inside-region. The number of force curves $N$ is displayed in the figure. Top row and bottom row are showing the exact same data in different combinations for comparison.

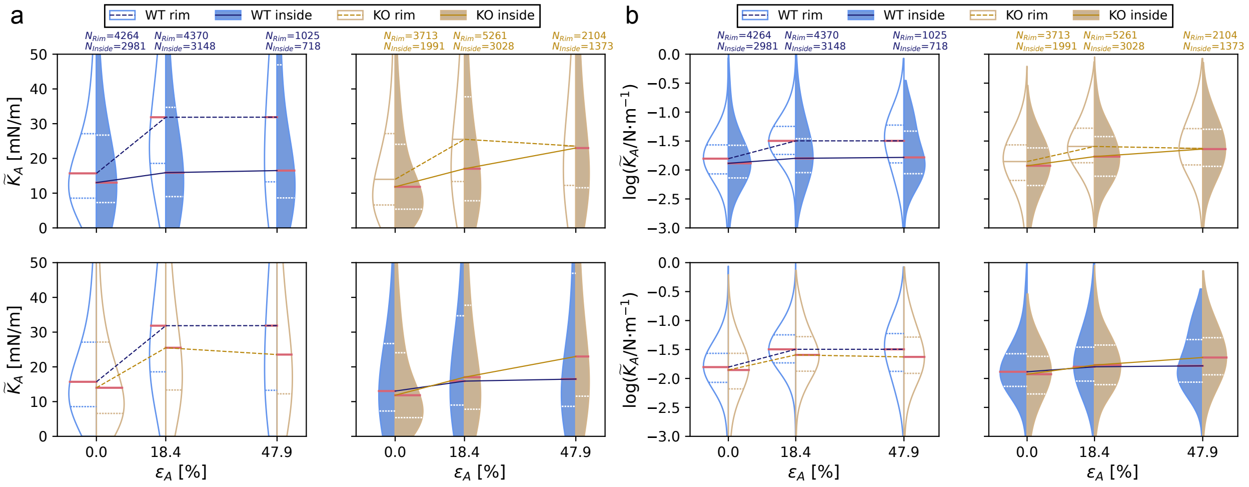


Figure S12: Area compressibility modulus $\widetilde{K}_A$ of cells under strain $\varepsilon_A$. Data for WT cells are shown in blue, data for KO cells in yellow. Filled violins correspond to the inside-region of the cells, open violins with colored outlines to the rim-region. The lines on the violins represent the median (solid pink line), the upper and lower quartiles (dashed lines). The medians are connected via lines: dashed lines for the rim-region and solid lines for the inside-region. The number of force curves $N$ is displayed in the figure. Top row and bottom row are showing the exact same data in different combinations for comparison. (a) Area compressibility modulus $\widetilde{K}_A$. (b) The decadal logarithm of the same data as in (a).

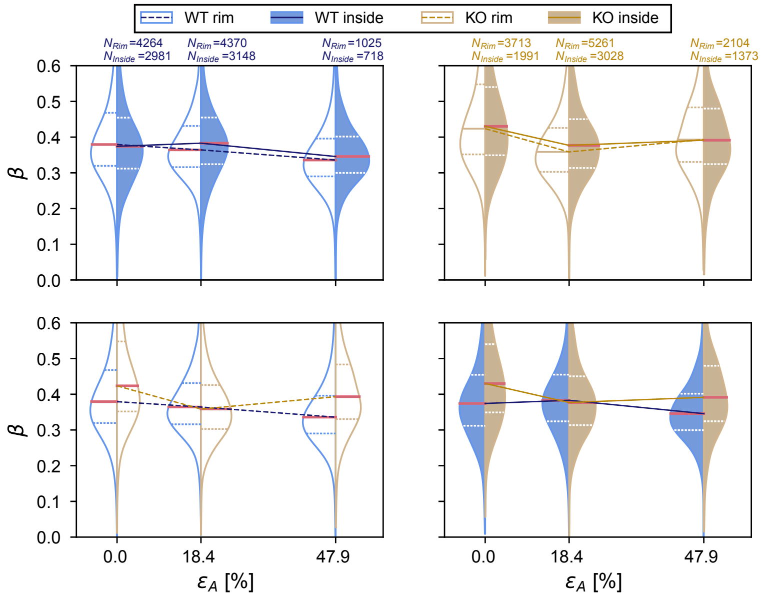


Figure S13: Fluidity $\beta$ of cells under strain $\varepsilon_A$. Data for WT cells are shown in blue, data for KO cells in yellow. Filled violins correspond to the inside-region of the cells, open violins with colored outlines to the rim-region. The lines on the violins represent the median (solid pink line), the upper and lower quartiles (dashed lines). The medians are connected via lines: dashed lines for the rim-region and solid lines for the inside-region. The number of force curves $N$ is displayed in the figure. Top row and bottom row are showing the exact same data in different combinations for comparison.

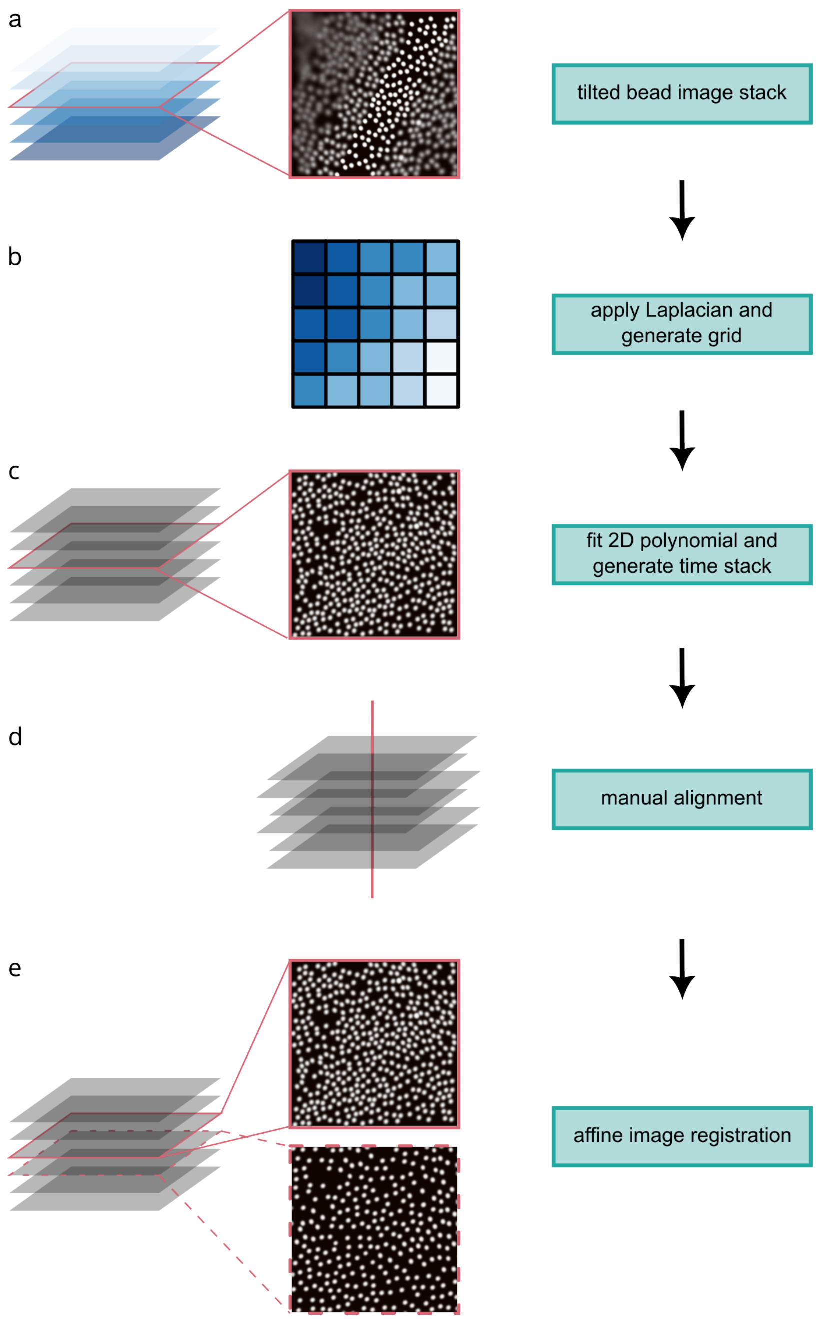


Figure S14: Workflow for the characterization of the PDMS devices in the equibiaxial stretcher. (a) Fluorescence image stack (different $z$-positions) of the beads in the slightly tilted membrane. (b) To compute the in-focus regions of the stack, a Laplacian is calculated for each frame and the indices of the frames in the stack where the beads are in focus are saved on a coarse grid. (c) A 2D polynomial is fitted to the grid and the original stack is interpolated to this fit to create a tilt-corrected image. By repeating this procedure for all image stacks with increasing strain, we generate a time stack with one tilt-corrected image per strain. (d) The images in the time-stack are manually aligned at the center of the image. (e) The images in the time stack are registered using an affine transformation to determine the strain.

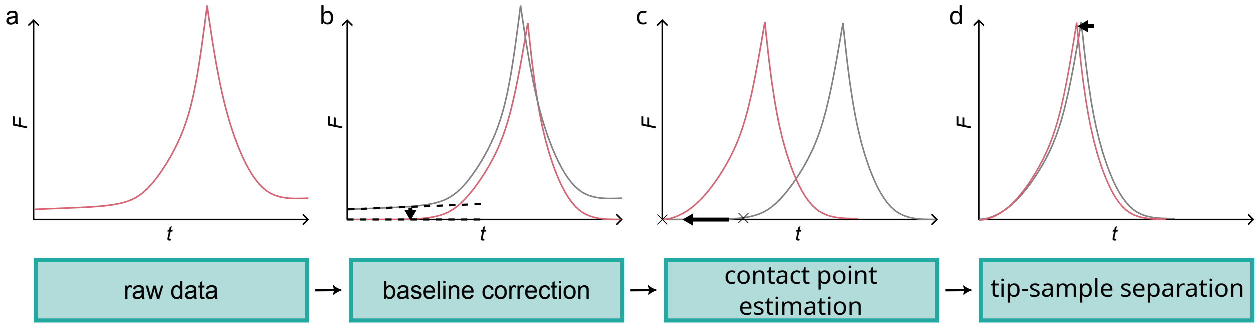


Figure S15: Schematic of the pre-processing of the force-time curves. (a) The baseline of the original force curve is (b) tilt and offset corrected by subtracting the baseline from the curve. (c) The contact point is estimated using the nanite-function "deviation from baseline" and the curve is shifted such that the contact point lies at time point 0. (d) Finally, a tip-sample separation is applied to separate the distance of the cantilever moved towards the sample from the deflection of the cantilever in the opposite direction.
